# Structural investigation suggests Elongation Factor-G1 is the *bona fide* canonical translation factor for ribosome recycling in mycobacteria

**DOI:** 10.64898/2026.05.25.727771

**Authors:** Ankit Dhur, Priya Baid, Krishnamoorthi Srinivasan, Jayati Sengupta

## Abstract

Bacterial elongation factor G (EF-G) facilitates mRNA and tRNA translocation during peptide elongation as well as promotes splitting of ribosomal subunits during ribosome recycling in coordination with ribosome recycling factor (RRF). Two homologs of EF-G (EF-G1 and EF-G2) have been identified in some bacterial species, including mycobacteria and in mitochondria. We have previously reported that mycobacterial EF-G2 functions as a stress-specific factor. Here, we demonstrate that EF-G1 functions as the canonical elongation factor responsible for ribosome recycling in mycobacteria. In addition, our structural analyses explain why mycobacterial EF-G2 lacks recycling activity, in contrast to its mitochondrial counterpart. We report cryo-EM structures of the *M. smegmatis* 70S ribosome and 50S subunit in complex with RRF and/or EF-G1 at 3-4 Å resolution, elucidating the molecular architecture of intermediate structures and mechanistic basis of ribosome recycling. Notably, GTP hydrolysis on EF-G1 is not required for ribosomal disassembly in mycobacteria, unlike in *E. coli*.

## Introduction

The elongation factor G (EF-G), a universally conserved translation factor belonging to the translation-associated GTPase family, serves dual functional roles in translation (1,2). It catalyses translocation during peptide elongation and mediates ribosomal disassembly during ribosome recycling in concert with the ribosomal recycling factor (RRF) (3–5). Following peptide bond formation, EF-G, in the presence of GTP, binds the pre-translocation state ribosome and catalyses the translocation of tRNAs from the A and P sites to the P and E sites, respectively, coupled with the shifting of the mRNA by one codon (4,6,7). In bacteria, at the end of a translation cycle, after the release of the polypeptide chain, deacylated tRNA and mRNA remain bound to the 70S ribosome (8,9). This complex (called the post-termination complex, PoTC) dissociates into the large 50S subunit and the small 30S ribosomal subunit, allowing them to participate in the next round of translation (10). The ribosome recycling factor (RRF) binds to the post-termination complex (PoTC), followed by the binding of the elongation factor G (EF-G), and after subsequent GTP hydrolysis, the PoTC is dissociated into the ribosomal subunits together with the release of deacylated tRNA and mRNA (8,9,11–14).

The structure of RRF consists of two domains: domain I is a long triple α-helix bundle, and domain II is composed of a smaller α/β domain. The two domains are linked through flexible hinges, allowing domain II to rotate freely around the long axis of domain I (15–17) (Fig. 1A). Different biochemical and structural studies have revealed that domain I of RRF binds across the A and P sites of the 50S ribosomal subunit, while domain II is flexible and is involved in bringing about conformational changes in the inter-subunit bridge B2a/d, thereby facilitating the dissociation of the ribosomal subunits in the presence of EF-G (18–20). Sequence comparison reveals that RRFs share approximately 85% identity among mycobacterial species and display around 40% sequence identity with RRF from *E.coli* (Supplementary Fig. 1A and C). The RRF of human mitochondria has an extra N-terminal extension (NTE), which the mycobacterial RRF lacks (21).

**Figure. 1:**
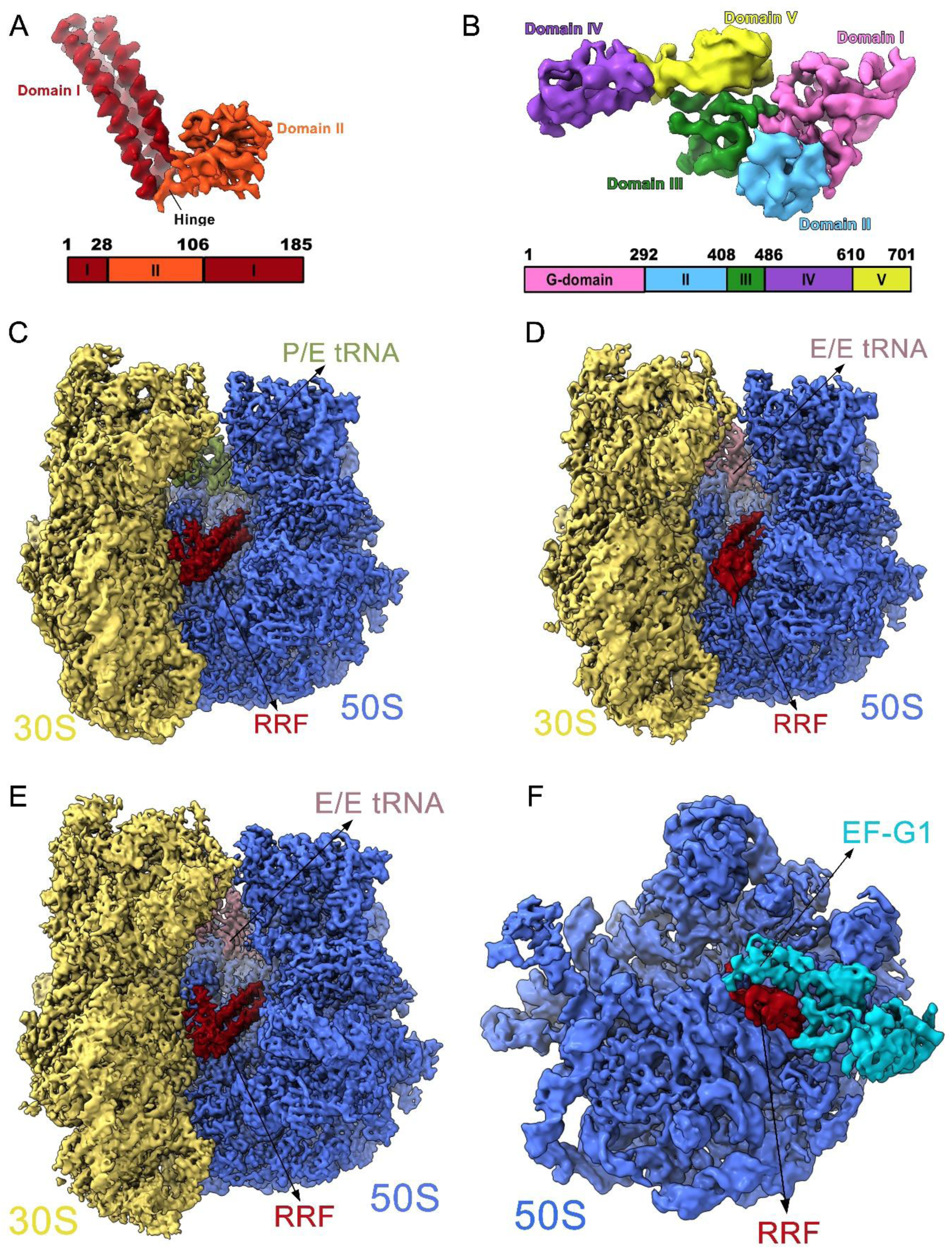
Cryo-EM maps of the intermediate states of the mycobacterial ribosome recycling process. (A) Extracted density of MsmRRF, domain I (red) and domain II (orange). (B) Extracted density of MsmEF-G1, domain I (pink), domain II (cyan), domain III (green), domain IV (purple) and domain V (yellow). (C) Cryo-EM map showing MsmRRF and P/E tRNA bound to the 70S ribosome (30S subunit ratcheted) with a resolution of 3.6 Å (Map 1). (D) Cryo-EM map showing MsmRRF and E/E tRNA bound to the 70S ribosome (30S subunit head swivelled) with a resolution of 3.8 Å (Map 2). (E) Cryo-EM map showing MsmRRF and E/E tRNA bound to the 70S ribosome (30S subunit head swivelled) with a resolution of 3.4 Å (Map 3). (F) Cryo-EM map showing MsmRRF and MsmEF-G1 bound to the 50S subunit with a resolution of 4.2 Å (Map 4). The 50S ribosomal subunit (dark blue), 30S ribosomal subunit (yellow), MsmRRF (red), MsmEF-G1 (cyan), P/E tRNA (light green) and E/E tRNA (light pink).

Most eubacteria harbour a single form of EF-G, which is involved in both translation elongation and ribosome recycling (1,2,8,9). Several bacteria, such as *Thermus thermophilus*, *Borrelia burgdorferi, Bacteroides thetaiotaomicron, and Pseudomonas aeruginosa,* harbour two homologs of EF-G, termed EF-G1 and EF-G2 (22–24). In *T. thermophilus,* EF-G1 participates in both translational elongation and ribosome recycling. *T. thermophilus* EF-G2 has been shown to possess both ribosome-dependent and intrinsic GTPase activity, along with translocation activity (25). However, its involvement in the ribosome recycling process remains unclear. In *B. thetaiotaomicron*, EF-G2 is known to play a role in slow translocation (24). In *B. burgdorferi*, EF-G1 is involved in the translational elongation process, while EF-G2 contributes to ribosome recycling (26). A similar division of labour between EF-G1A and EF-G1B has been observed in *P. aeruginosa* (22). Interestingly, two EF-G homologs have also been found in human mitochondria: EF-G1mt and EF-G2mt. EF-G1mt participates in elongation, while EF-G2mt is responsible for ribosome recycling (21,27,28).

Mycobacteria also harbour two EF-G paralogues, EF-G1 (sharing 60% amino acid identity with the *E. coli* EF-G), and EF-G2, which shares 31% amino acid identity with *E. coli* EF-G. *M. smegmatis* (Msm) EF-G1 and EF-G2 share 31% amino acid identity and 51% amino acid similarity with each other (29) (Supplementary Fig. 1D-F). However, the functional roles of the two EF-Gs in mycobacterial ribosome recycling remained unknown. In a recent study on mycobacterial EF-Gs, we have shown that EF-G2 exhibits minimal ribosome-dependent GTPase activity and is incapable of performing ribosome recycling. Instead, it has a regulatory role in translation under stress conditions(30).

This study aims to uncover the structural basis of the ribosome recycling process in mycobacteria by determining high-resolution cryo-EM structures of the ribosome recycling complexes. Our findings suggest that, unlike the division of labour reported for mitochondrial EF-Gs, in mycobacteria, EF-G1 serves as the canonical translation factor, whereas EF-G2 carries out stress-associated roles. Here, we present seven cryo-EM maps (resolutions ranging 3-4 Å), comprising six *M. smegmatis* 70S ribosome (Msm70S) and one 50S subunit (Msm50S) complexes with MsmRRF and/or MsmEF-G1 elucidating the mechanism of ribosome recycling at the molecular level. Of note, it was found that GTP hydrolysis on EF-G1 is not required for the ribosomal disassembly in mycobacteria.

Our analyses further clarified the structural features that enable MsmEF-G1 to mediate ribosome recycling in cooperation with MsmRRF, while rendering MsmEF-G2 incapable of performing this function. Interestingly, one of the structures described here is the tRNA-bound 70S ribosome in complex with MsmEF-G1, representing a well-characterised tRNA translocation intermediate in which the 30S subunit adopts a ratcheted conformation, as reported by Zhou et al. (31), indicating that EF-G1 also participates in tRNA translocation.

## Materials and methods

### PCR amplification and cloning of genes

The genes encoding *M.smegmatis* RRF and EF-G1 were PCR amplified from genomic DNA using specific primers given below and subcloned into kanamycin-resistant pET28a DNA vector (Novagen) containing requisite restriction sites.

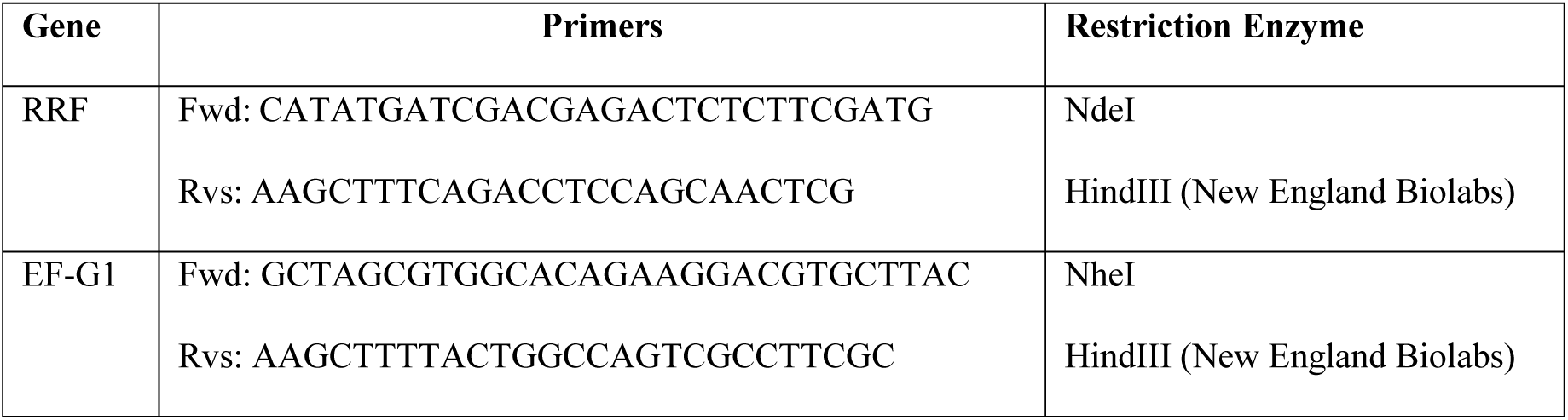

The *E.coli* infC gene (IF3) cloned in ampicillin resistant pET19a vector was purchased from Bio Bharati Life Science.

### Expression and purification of recombinant proteins

The positively cloned vectors were transformed into *E. coli* BL21 (DE3) for expression. For *M.smegmatis* RRF, the culture was grown till log phase (OD_600_= 0.6) and was induced with 0.25mM isopropyl-β-D-thiogalactopyranoside (IPTG) and incubated at 37 °C for 2 hr 30 minutes. Cells were harvested and lysed in lysis buffer (50 mM Tris-HCl, pH 7.6, 150 mM NaCl, 5% glycerol, 2 mM 2-mercaptoethanol, 1 mM PMSF and 2mM Imidazole) containing lysozyme by ultrasonication. Ni-NTA chromatography was used to purify the 6x His-tagged recombinant protein. After centrifugation of cell lysates at 17000 rcf for 45 minutes at 4 ⁰C, the supernatant was loaded onto a column containing Ni-NTA resins (GE Healthcare) washed with wash buffer (50 mM Tris-HCl, pH 7.6, 150 mM NaCl, 5% glycerol and 35 mM imidazole) and eluted with elution buffer (50 mM Tris-HCl, pH 7.6, 150 mM NaCl, 5% glycerol and 150 mM imidazole). The protein enriched fractions were pooled together, and buffer-exchanged with storage buffer (20mM Tris-HCl, pH 7.6,50mM NH_4_Cl, 10mM Mg-acetate, 3mM 2-mercaptoethanol and 5% glycerol) and concentrated by using Amicon Ultra-4 Centrifugal Units (Merck) and the aliquots were stored in -80 °C. For *M.smegmatis* EF-G1 the log phase culture (OD_600_= 0.6) was induced with 0.1mM isopropyl-β-D-thiogalactopyranoside (IPTG) and incubated at 37 °C for 4 hours. The same purification protocol was followed for MsmEF-G1, but with changes in the buffer composition.

Lysis buffer: 50mM HEPES-KOH, PH 7.6, 500mM NaCl, 1mM PMSF, 5mM Imidazole, 1mM DTT and 8% glycerol.

Wash buffer: 50mM HEPES-KOH, PH 7.6, 500mM NaCl, 40mM Imidazole, and 8% glycerol. Elution buffer: 50mM HEPES-KOH, PH 7.6, 500mM NaCl, 150mM Imidazole, and 8% glycerol.

Storage Buffer: 20 mM HEPES-KOH; pH 7.6, 50 mM NaCl, 50 mM KCl, 5 mM Mg-acetate, 1 mM DTT, 8% glycerol.

The purified protein was concentrated by using Amicon Ultra-4 Centrifugal Units (Merck) in storage buffer, and the aliquots were stored in -80 °C.

*E.coli* IF3 was purified using the same protocol as *M.smegmatis* RRF but with some changes in the buffer composition.

Lysis Buffer: 20 mM Tris-HCl, pH 7.5, 250 mM NaCl, 10% glycerol, 2 mM 2-mercaptoethanol and 1 mM PMSF.

Wash Buffer: 20 mM Tris-HCl, pH 7.5, 250 mM NaCl, 5% glycerol, 2 mM 2-mercaptoethanol, 30mM Imidazole and 1 mM PMSF.

Elution Buffer: 20 mM Tris-HCl, pH 7.5, 250 mM NaCl, 5% glycerol, 2 mM 2-mercaptoethanol, 150 mM Imidazole and 1 mM PMSF.

Storage Buffer: 20 mM Tris-HCl, pH 7.5, 250 mM NaCl, 5% glycerol and 3 mM 2-mercaptoethanol.

### Purification of ribosomes from *Mycobacterium smegmatis*

The 70S ribosome was isolated and purified based on the protocol by Baid. P et al(30). *Mycobacterium smegmatis* mc^2^155 cells were grown till late log phase (OD_600_= 1.1) and then harvested by centrifugation. The cell pellet was washed thrice with Buffer A (20 mM Tris–HCl; pH-7.6, 100 mM NH_4_Cl, 10 mM Mg-acetate and 5 mM β-ME) and again resuspended in the same buffer and lysed by sonication. The cell debris was removed by centrifugation, and the supernatant was subsequently subjected to ultra-centrifugation using a swing-out rotor Thermo Scientific™ AH-629 (154,000g, 4 °C, 2 hours 30 minutes). The crude ribosome pellet was dissolved in 500 μL of buffer ‘B’ (20 mM Tris–HCl; pH 7.6, 30 mM NH_4_Cl, 10 mM Mg-acetate and 5 mM β-ME) and stored overnight at 4 °C. The next day, the crude ribosome pellet was salt-washed and homogenised in buffer ‘C’ (20 mM Tris–HCl; pH 7.6, 1 M NH_4_Cl, 10 mM Mg-acetate and 5 mM β-ME) for approximately 2 hours. The homogenised ribosome preparation was centrifuged, and the supernatant was ultra-centrifuged using a swing-out rotor Thermo Scientific™ AH-629 (154,000g, 4 °C, 2 hours 30 minutes). The pellet was dissolved in Buffer B and loaded on top of a 10%–40% linear sucrose gradient (made with Buffer B) and ultra-centrifuged using swing-out rotor Thermo Scientific™ AH-629 (154,000g, 4 °C, 4 hours 30 minutes). Fractions were collected manually from bottom to top from the sucrose gradient using a peristaltic pump. The absorbance at 260 nm was measured for all the fractions, and the fractions corresponding to the 70S ribosome were pooled together and concentrated using Buffer B using Amicon Ultra-4 Centrifugal Units (Merck) and stored in -80 °C.

### Site-directed mutagenesis (SDM) of M. smegmatis EF-G1

Point mutations were introduced into the M. smegmatis EF-G1 gene cloned in pET28a by site-directed mutagenesis using mutation-specific primer pairs (listed below).

Msm G1_R62Q FWD: 5’- GGAGCAGGAGCAGGGTATCACCATC - 3’

Msm G1_R62Q RVS: 5’- GATGGTGATACCCTGCTCCTGCTCC - 3’

Msm G1_H88Y FWD: 5’- CACCCCCGGCTACGTCGACTTC - 3’

Msm G1_H88Y RVS: 5’- GAAGTCGACGTAGCCGGGGGTG - 3’

Following PCR amplification, the reaction mixture was treated with DpnI (New England Biolabs, USA) to selectively digest the parental methylated plasmid DNA. The resulting amplified products were transformed into ultra-competent E. coli DH5α cells and plated on LB agar supplemented with kanamycin, followed by incubation at 37 °C for 14–16 hours. Individual colonies were selected for plasmid isolation, and the presence of the desired mutations was verified by DNA sequencing. Sequence-verified plasmids carrying EF-G1 point mutations were subsequently transformed into *E. coli* BL21 (DE3) pLysS cells for mutant protein expression.

Mutated proteins were expressed and purified in the same way as the wild-type protein of MsmEF-G1, as mentioned above.

### Ribosome-dependent GTPase assay of wild-type and mutant mycobacterial EF-G1

Ribosome-dependent GTPase activity of EF-Gs was measured using a malachite green phosphate assay kit, which detects inorganic phosphate released during GTP hydrolysis through the formation of a green-colored complex under acidic conditions. Assays were performed using wild-type and mutant EF-G1 proteins. Reactions were carried out in GTPase buffer containing 20 mM Tris–HCl (pH 7.6), 40 mM NH₄Cl, 30 mM KCl, 10 mM Mg-acetate, and 1 mM DTT. The reaction mixture consisted of 100 nM *M. smegmatis* 70S ribosomes, 1 µM EF-G protein, and 100 µM GTP, and was incubated at 37 °C. Aliquots (100 µl) were withdrawn at 10-min intervals over a 60-min period and immediately mixed with malachite green reagent. Following a 2-min incubation at room temperature, absorbance at 630 nm was recorded, and phosphate release was plotted as a function of time.

### Sample preparation for Cryo-EM and data collection

The complex of *M.smegmatis* RRF, EF-G1 with 70S ribosome was prepared in Binding Buffer (50 mM HEPES–KOH; pH-7.6, 70 mM NH4Cl, 30 mM KCl, 7 mM Mg-acetate and 1 mM DTT) in the presence of non-hydrolysable GTP analogue GMP-PNP. The ribosome (0.7uM) was pre-treated with 700uM of puromycin. To this puromycin-treated ribosome, a 30-fold excess of RRF relative to the ribosome concentration was added and incubated at 32 °C for 15 minutes. To this mixture, 10-fold excess of EF-G1, 2-fold excess of IF3 (both relative to the concentration of the ribosome) along with a 100-fold excess of GMP-PNP relative to the EF-G1 concentration were added. Then the mixture was incubated at 32 °C for 15 minutes and kept on ice. This complex was loaded on a sucrose cushion (25% w/v sucrose in binding buffer) and ultra-centrifuged using the T-890 rotor (Thermo Scientific) (196 000 g, 4 °C, 6 hours). The pellet was resuspended in binding buffer, and the protein-bound ribosome complex was diluted to a final concentration of 200 nM and kept on ice for the grid preparation. We used Quantifoil R2/2 carbon-coated holey grid, which was glow-discharged. The grids were prepared with Vitrobot Mark IV (FEI, Hillsboro, OR, USA). High-resolution data were acquired on a 300-kV FEI Titan Krios electron microscope (Thermo Fisher Scientific) at the National Center for Biological Science (NCBS)-National Electron Cryo-Microscopy Facility, Bangalore Life Science Cluster (BLiSc), fitted with Falcon III direct electron detector. A total of 3,899 movie stacks were recorded, each consisting of 20 frames, with an image pixel size of 1.07 Å and an electron dose of 1.04 e⁻/Å² per frame.

### Cryo-EM data processing

The data was processed using Relion-3.0.8 and Relion-3.1. Dose-induced motion correction was done using Relion’s own implementation, MotionCor2, and CTF estimation of the motion-corrected movies was carried out using CTFFIND 4.0. Micrographs with good CTF values were selected, and 644,542 particles were autopicked from them. These autopicked particles were subjected to iterative rounds of 2D classification, and the best 2D classes were chosen corresponding to 264,233 centred and cleaned particles. These particles underwent multiple rounds of reference-based 3D classification. The 70S and 50S ribosome populations were separated. The 50S ribosome class was further subjected to 3D classification, yielding a subset containing RRF and EF-G1 bound to the 50S subunit (15,686 particles). This class underwent CTF refinement and particle polishing and was refined to a resolution of 6.5Å (FSC criteria 0.143) and postprocessed resolution of 4.4Å (Map 4). Similarly, the 70S ribosome class was subjected to 3D classification, yielding three distinct protein and tRNA-bound 70S ribosome classes. These classes further underwent focused 3D classification using an appropriate mask. After the classification, the first 70S ribosome class yielded two distinct subclasses: a RRF-bound 70S ribosome (61,327 particles) and another, a P-site tRNA-bound 70S ribosome (36,773 particles). Both classes underwent CTF refinement and particle polishing, and after 3D refinement, the RRF-bound 70S ribosome class had a resolution of 3.7Å (FSC criteria 0.143) and postprocessed resolution of 3.1Å, while the P tRNA-bound 70S ribosome class had a resolution of 3.9Å (FSC criteria 0.143) and postprocessed resolution of 3.2Å. The second 70S ribosome class produced three subclasses: two RRF-bound 70S ribosomes with an E/E state tRNA (Maps 2 and 3), having 18,050 and 32,720 particles respectively and one EF-G1-bound 70S ribosome with 13,826 particles. Map 2, after 3D refinement, achieved a resolution of 5.1Å (FSC criteria 0.143) and postprocessed resolution of 3.8Å. Map 3, after refinement, had a resolution of 4.2Å and postprocessed resolution of 3.4Å and the 70S-EF-G1 map had a resolution of 5.1Å (3D refined) and 3.7Å (postprocessed). The third 70S ribosome class did not yield any additional subclass after focused 3D classification. This class was 3D refined and achieved a resolution of 4.5Å and a postprocessed resolution of 3.6Å (Map 1). The local resolution of all the maps was estimated using ResMap (Supplementary Fig. 2 and 3).

### Model building, validation and structural analysis

The refined maps were sharpened using a model-based local sharpening tool in PHENIX (version 1.20.1-4487), and it was used for the preparation of the structural illustrations. The model coordinates of the *M.smegmatis* (*Msm*)70S ribosome (PDB-9K0Z) and 50S ribosome (PDB-7Y41) were used as a starting model for model building. For MsmRRF and EF-G1 an alpha fold generated model coordinate was used as the initial template. All the model coordinates were initially fitted into the maps using CHIMERA. Further, different regions of the model were manually fitted using WinCoot (version 0.9.8.1) and refined using Phenix Real Space Refinement iteratively. The models were validated using MolProbity in PHENIX. The illustrations were made using Chimera, ChimeraX and PyMOL (Schrödinger, LLC.).

## Results and Discussion

### Cryo-EM captured intermediate states of the mycobacterial ribosome recycling process

We have prepared a complex of the mycobacterial 70S ribosome with MsmRRF and MsmEF-G1 in the presence of GMPPNP, a non-hydrolysable GTP analogue. The use of GMPPNP inhibits the GTP hydrolysis-dependent release of EF-G1 from the ribosome, thereby stabilising intermediate states that would otherwise be difficult to capture due to the rapid transition of GTP hydrolysis-driven states. This provides us with a detailed structural characterisation of the recycling process. We have included *E.coli* IF3 while preparing the complex, where its role was to prevent the re-association of dissociated ribosomal subunits (12,32–34).

We have processed the cryo-EM dataset in RELION(35), and after multiple rounds of 2D and 3D classification, obtained seven distinct classes, six 70S ribosome complexes and one 50S subunit complex (Supplementary Fig. 2 and 3). Among these, four classes represent distinct intermediate states of the ribosome recycling process. Map 1, resolved at 3.6 Å, represents a distinct structural intermediate in the ribosome recycling process, in which the orientation of the RRF domain II points towards the 30S subunit and interacts with ribosomal protein bS12 (Fig. 1C and 2A). When compared with the non-rotated 70S-P site tRNA class, the 30S subunit of Map 1 shows ratchet-like motion, characterised by relative rotational movement between the head and body of the subunit(36) (Fig. 2C). Further, there is a shifting in the position of the L1 stalk on the 50S ribosomal subunit. In this map, the tRNA adopts a hybrid P/E configuration, where the anticodon arm remains positioned between the P and E sites on the 30S subunit, while the CCA end extends into the E site on the 50S subunit (Fig. 2E). This intermediate conformation of the tRNA closely resembles the P/E state, reported by C. E. Carbone *et al* in PDB 7SSN(6).

**Figure. 2:**
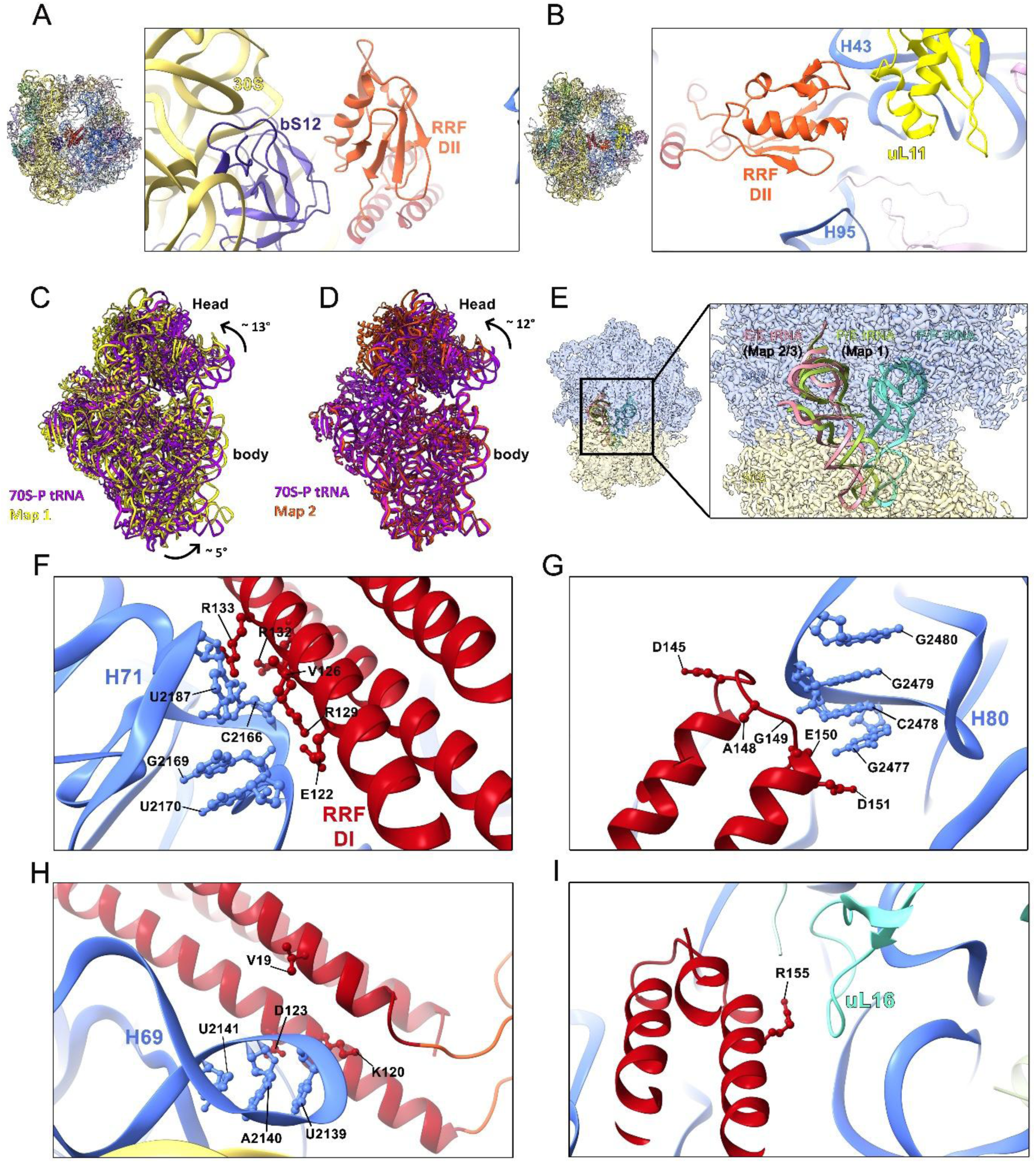
Structural features of MsmRRF-bound recycling intermediate states of the mycobacterial 70S ribosome. (A) Interactions of the 30S (yellow) ribosomal protein bS12 (blue) with MsmRRF domain II (DII) (orange) in Map 1 are displayed. (B) Interactions of MsmRRF domain II (orange) with 50S (blue) ribosomal protein uL11 (yellow), H43 and H95 in Map 2 are displayed. (C, D) The conformational dynamics of the ribosomal 30S subunit. Superposition of the 30S subunits in the structures 70S-P tRNA and Map 1 (C), showing the ratcheted motion of the 30S subunit of Map 1 and 70S-P tRNA and Map 2 (D), showing head swivelling in the 30S subunit of Map 2. The 30S subunit of 70S-P tRNA (purple), Map 1 (yellow) and Map 2 (orange). (E) The tRNA dynamics during the recycling process. The P/P tRNA (cyan), P/E tRNA (light green) and E/E tRNA (light pink). (F-I) Snapshots of the molecular interaction between MsmRRF and H71, H80, H69 and the ribosomal protein uL16 in Map 2. The 23S rRNA (dark blue), uL11 (yellow), MsmRRF Domain I (D1) (red) and uL16 (cyan).

The second structure (Map 2) has a resolution of 3.8 Å, shows RRF bound to the 70S ribosome, with domain II pointing towards the 50S subunit and has the presence of a tRNA occupying a different position than Map 1 (Fig. 1D and 2B). The domain II of RRF is oriented towards the 50S subunit in proximity to helix 43 (H43), H95 (part of the SRL) of the 23S rRNA and the ribosomal protein uL11 (Fig. 2B), while the tRNA occupies the E site on the ribosome. From the structural analysis, we observe that the domain I of RRF is quite rigid and has a stable conformation in contrast to the more flexible domain II. This stability is due to the multiple interactions of its domain I amino acid residues with the ribosome. The amino acid residues Glu122, Val126, Arg129, Arg132 and Arg133 interact with the rRNA (nt C2166, G2169, U2170, U2187) of the H71 (Fig. 2F) while Glu147, Ala148, Gly149 and Glu150 make contact with the nucleotides (nt 2478-2480) of H80 of the 23S rRNA (Fig. 2G). H69 nucleotide residues (nt 2134, 2140-2142) also make contacts with the domain I of RRF (Fig. 2H). Additionally, the residue Arg155 of the RRF domain I has an interaction with the loop of the ribosomal protein L16 (Fig. 2I). A pronounced 30S subunit head rotation (swivelling) is also evident in Map 2 when compared with the non-rotated 70S ribosome class containing P site tRNA (Fig. 2D).

The third 70S structure (Map 3) resolved at 3.4 Å, shows that the overall orientation of RRF, including its position of domain II, remains comparable to that observed in Map 1; however, the position of the tRNA is more towards the E site, matching the conformation observed in Map 2 (E/E state) (Fig. 1E and 2E). Likewise, the 30S subunit also adopts a configuration similar to that in Map 2 (Fig. 2D).

The fourth recycling intermediate (Map 4) corresponds to the 50S subunit and is resolved at a lower resolution of 4.4 Å. The map shows both RRF and EF-G1 bound to the 50S subunit (Fig. 1F). Despite having a lower resolution, all five domains of EF-G1 were well-defined and could be visualised interacting with the RRF (Fig. 1B and F). Across all four classes representing recycling intermediates, domain I of RRF was consistently well resolved, suggesting it adopts a stable conformation following ribosome binding. In contrast, domain II appeared more dynamic, which is evident from its comparatively lower local resolution (Supplementary Fig. 3). Structurally, the RRF from *M. smegmatis* adopts the characteristic L-shaped structure commonly observed in other eubacterial species (15–17,37)(Fig. 1A).

Earlier studies reported some intermediates that we also observed, but resolved at much lower resolution (∼7-16 Å). In the time-resolved study by Z. Fu *et al*., one intermediate closely corresponds to one of our obtained class (non-ratcheted 70S-RRF)(18) (Supplementary Fig. 4A). This structure, termed NR-PostTC.RRF140 (resolution 16 Å) features RRF bound to a non-ratcheted 70S ribosome, with domain II of RRF positioned toward and interacting with the stalk base of the 50S subunit. In this structure, they have observed a shift of RRF domain I toward the peptidyl transferase centre. Our high-resolution maps consistently show domain I occupying the same position across all classes. In our 70S-RRF structure, we have observed interaction of H69 residues with RRF domain I, along with H71 and H80 of the 23S rRNA, similar to that observed in our Map 2, which were not distinct at their resolution (Fig. 2F-I).

Similarly, in a related study by T. Yokoyama *et al*., reported a recycling intermediate in which RRF is bound to the 70S ribosome with its domain II towards the 50S subunit and a bound tRNA at a resolution of 11Å, which closely corresponds to our Map 2 (resolution 3.8 Å)(19). Furthermore, they have reported an interaction between the elbow region of RRF and protein bS12; in contrast, our higher-resolution map does not show any direct contact between RRF and the 30S subunit. In agreement with their findings, we also observe interactions between RRF and helices H69, H71, and H95, in addition, our maps, resolved at higher resolution, reveal further contacts involving H80 and ribosomal protein uL16, which were not reported by them (Fig. 2G and I). They have also observed a displacement of H43 in the 23S rRNA upon RRF binding. However, when we compared our Map 2 with the 70S-P site tRNA-bound class, we did not observe any corresponding shift in H43 (Supplementary Fig. 4C).

The remaining two classes comprise a non-ratcheted 70S ribosome with a P site-bound tRNA, and an EF-G1 bound to a ratcheted 70S ribosome with a bound tRNA. The resolutions of the three classes were 3.2 Å, 3.1 Å, and 3.7 Å, respectively (Supplementary Fig. 2 and 3).

### Mycobacterial EF-G1 has a conserved G domain architecture capable for ribosome-dependent GTPase activity

The G domain (Domain I) of EF-G is responsible for GTP binding and hydrolysis(38–42). It consists of five important conserved GTP-binding motifs (G1-G5) (41,42). Structural and amino acid sequence comparison of MsmEF-G1 with canonical bacterial EF-G shows that these motifs are highly conserved in MsmEF-G1 (29,43) (Supplementary Fig. 1B and Fig. 3A). The G1 motif (AHIDAGKT), which forms part of the P-loop, plays a central role in interacting with the α and β phosphates of GTP and proper positioning of the tri-phosphate moiety of the nucleotide (38,40). In MsmEF-G1, the P-loop residues His21, Ile22 and Asp23 interact with the highly conserved sarcin-ricin loop (SRL) of the 23S rRNA, which helps in the activation of GTP hydrolysis (Fig. 3B). In contrast, in MsmEF-G2, the amino acid stretch (Ala20-Ala24) is replaced by Gly30, Pro31, Ser32, Gly33 and Gly34, which results in the loss of key interactions (Fig. 3D). The G2 motif residues (ERGITI) constitute part of the switch I. In MsmEF-G1, the residues Arg62, Ile64, and Thr65 are very crucial for its GTPase activity, where Arg62 and Ile64 have critical interactions with the SRL(44). Notably, none of these residues is conserved in MsmEF-G2 (Fig. 3C). The G3 motif (DTPGH) forms part of switch II, in which the catalytic His88 forms a crucial interaction with the γ-phosphate of GTP (38,40,45). In MsmEF-G1, the catalytic His88 is precisely positioned towards the γ-phosphate of the nucleotide through a hydrophobic gate formed by Ile22 of the P-loop and Ile64 of switch I(40,43) (Fig. 3B). In MsmEF-G2, however, this catalytic His88 is replaced by Tyr98, which is oriented away from the γ-phosphate, abolishing its interaction with the nucleotide (Fig. 3D). The G4 motif (NKMD) forms crucial interactions with the guanine base and plays a major role in the specificity of the nucleotide base(40,43) (Fig. 3A). In this motif, the conserved Asn138 and Met140 of MsmEF-G1 are replaced by Thr148 and Leu150 in MsmEF-G2 (Fig. 3C). Finally, the G5 motif (GSA), which also contributes to the recognition of the guanine base, shows that none of the residues is conserved in MsmEF-G2 (Fig. 3C).

**Figure. 3:**
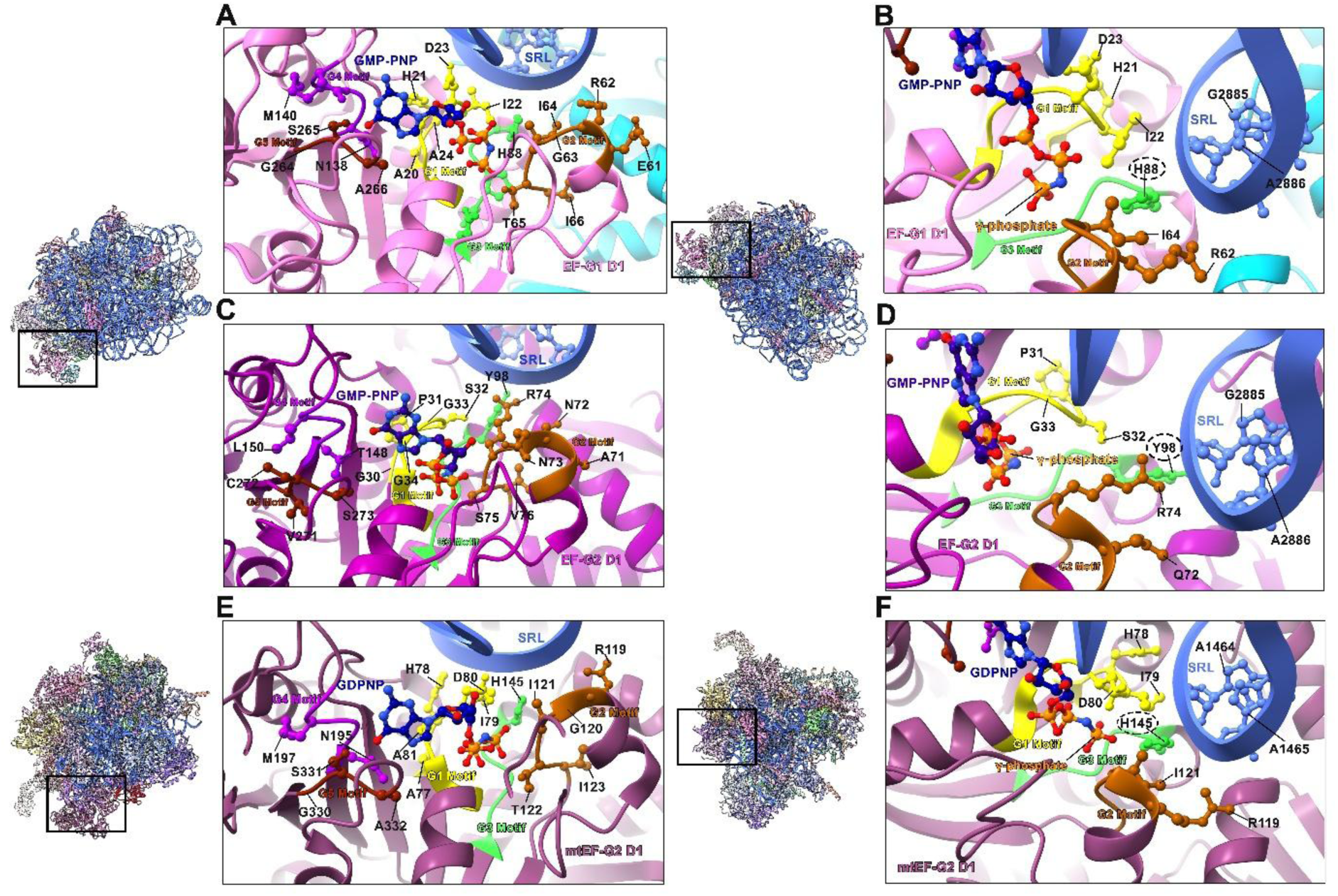
Comparison of the GTP-binding pocket of MsmEF-G1 and MsmEF-G2. (A,B) Close-up views of the GTP-binding motifs: G1 motif (yellow), G2 motif (brown), G3 motif (green), G4 motif (brick red) and G5 motif (purple) (A) and the SRL (blue) interacting residues (B) of MsmEF-G1 domain I (D1) (pink) with catalytic histidine H88 is seen to be oriented towards the γ-phosphate (orange) of GMP-PNP (dark blue). (C,D) Close-up views of the GTP-binding motifs and the SRL-interacting residues showing the altered amino acid residues in MsmEF-G2 domain I (D1) (magenta), specifically, the catalytic H88 is replaced by Y98 and is seen to be oriented away from the γ-phosphate. (E, F) Close-up views of the GTP-binding motifs and the SRL-interacting residues in mitochondrial EF-G2 domain I (mtEF-G2 D1) (dark pink), specifically, showing the conserved GTP-binding pocket, the residues interacting with the SRL and the catalytic H145 oriented towards the γ-phosphate (orange) of GDPNP (dark blue).

To further validate the role of switch region residues in GTP hydrolysis, we introduced targeted point mutations in MsmEF-G1. Specifically, we generated an R62Q substitution in switch I and an H88Y substitution in switch II region. The ribosome-dependent GTPase activities of these mutants were then assessed. Compared with the wild-type protein, the R62Q mutant exhibited a marked reduction in GTPase activity, whereas the H88Y mutant showed near complete loss of GTPase activity (Supplementary Fig. 5D). This observation thus proves the critical contribution of these conserved residues in MsmEF-G1 on GTP hydrolysis as observed in the case of other canonical EF-Gs.

It has been shown that human mitochondrial EF-G2 (mtEF-G2) participates in ribosome recycling in conjunction with human mitochondrial RRF (mtRRF)(21,27,28). The mtEF-G2 has ribosome-dependent GTPase activity, which is essential for the recycling process. Structural analysis of mtEF-G2 domain I shows that GTP-binding motifs (G1-G5) are well conserved (Supplementary Fig. 1B and 3E). The catalytic residue His145 is properly oriented towards the γ-phosphate of the bound nucleotide, supporting efficient GTP hydrolysis (Fig. 3F). In addition, the residues involved in crucial interactions with the SRL are also conserved in mtEF-G2 (Fig. 3F).

### EF-G1 serves as the canonical translation factor, in contrast to EF-G2 in mycobacteria

We have shown that in Mycobacterium, EF-G1 is capable of performing ribosome recycling, while EF-G2 is not(30). Previous reports on MsmEF-G2 have shown that it cannot effectively replace MsmEF-G1, as it fails to rescue the growth of an *E. coli* strain harbouring a temperature-sensitive allele of EF-G (fusA^ts^) at a non-permissive temperature (29). Moreover, the disruption of the MsmEF-G2 gene does not affect the growth kinetics of *M.smegmatis*. Recent cryo-EM structures from our laboratory elucidated the mechanistic basis of this functional divergence, showing that mycobacterial EF-G2 lacks ribosome-dependent GTPase activity, a crucial requirement for catalyzing these processes (30).

To understand what makes EF-G1 structurally and functionally compatible for ribosome recycling but not MsmEF-G2, we compared the ribosome-bound structures of EF-G1 and EF-G2, focusing on their domain organization and interaction with RRF. Structural comparison of the Msm50S–RRF–EF-G1 complex (Map 4) and the Msm50S–EF-G2 complex (PDB 9K10) (30) revealed a distinct difference in the position of domain I (G domain). While the EF-G1 domain I is closely positioned near the SRL, facilitating the interactions necessary for GTPase activation. In contrast, the MsmEF-G2 domain I is found to be shifted away from the SRL (Fig. 4A). Additionally, as mentioned previously, the SRL-interacting residues, as well as the residues crucial for the catalysing GTP hydrolysis in MsmEF-G1, are also not conserved in MsmEF-G2. These factors abolish efficient GTP hydrolysis by MsmEF-G2, whereas MsmEF-G1 is fully competent for this function.

**Figure. 4:**
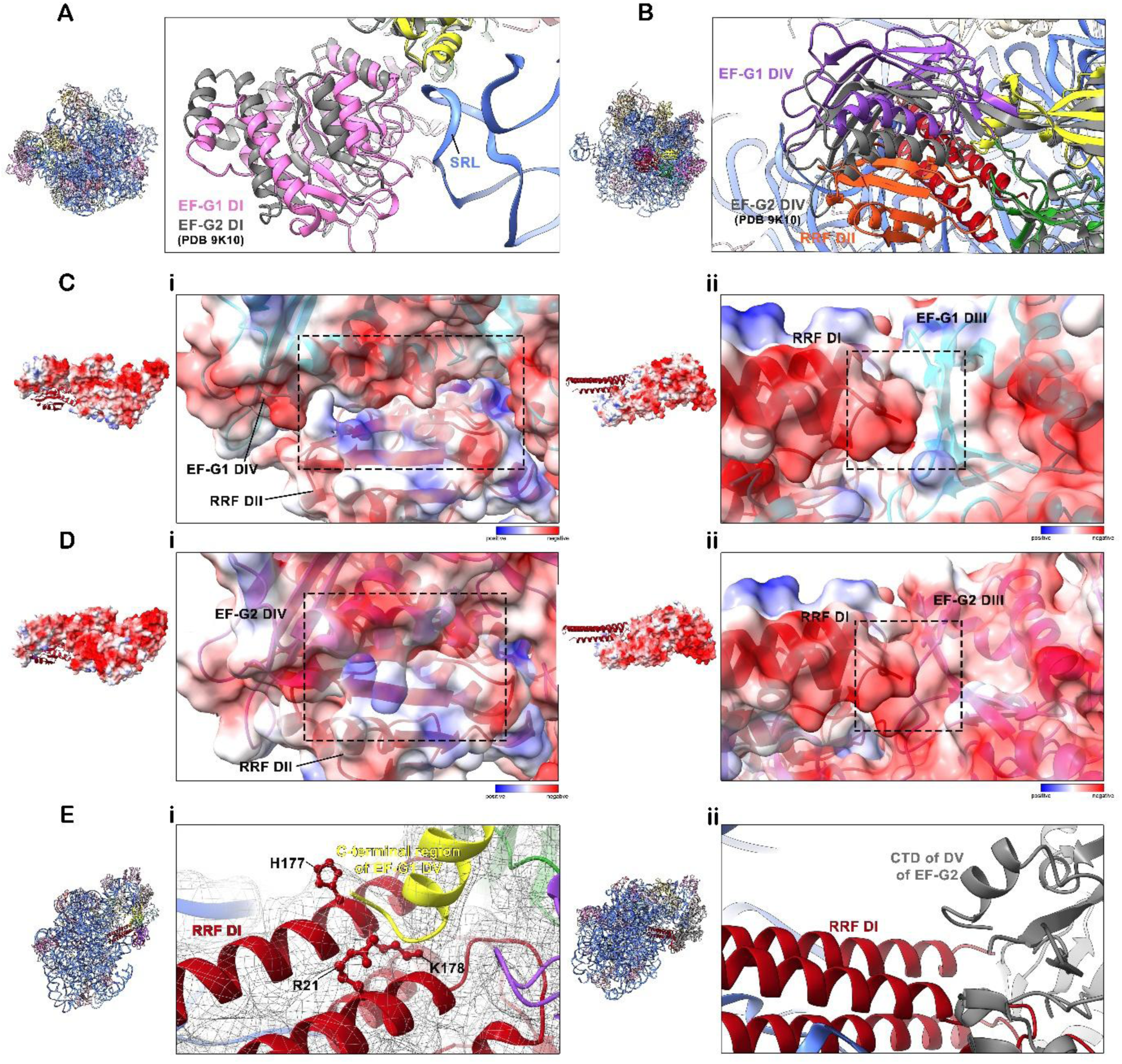
Comparison of the interaction of MsmEF-G1 and MsmEF-G2 with MsmRRF and the SRL in the 50S subunit complex. (A) Comparison of the position of MsmEF-G1 domain I (D1) (pink) with MsmEF-G2 domain I (D1) (PDB 9K10) (grey), showing that the latter is shifted away from the SRL (blue). (B) Comparison of the position of MsmEF-G1 domain IV (DIV) (purple) with Msm EF-G2 DIV (grey), showing the latter is shifted more towards the RRF domain II (DII) (orange), which will lead to a near steric clash. (C) Electrostatic surface representation of the Msm50S-EF-G1–RRF complex (Map 4). The boxed region shows the favourable charge compatibility between MsmRRF DII and MsmEF-G1 DIV (Ci) and MsmRRF DI and MsmEF-G1 DIII (Cii). (D) Electrostatic surface representation of the Msm50S-EF-G2–RRF complex (PDB 9K10). The boxed region shows the charge incompatibility between MsmRRF DII and MsmEF-G2 DIV (Di) and MsmRRF DI and MsmEF-G2 DIII (Dii). Negatively charged regions are shown in red and positively charged regions in blue. (E) Close-up views showing the interaction between MsmRRF domain I (D1) (red) and the C-terminal region of domain V (DV) of MsmEF-G1 (yellow) (Ei) and MsmEF-G2 (grey) (Eii), showing a productive interaction formed by the C-terminal region of DV of MsmEF-G1 with MsmRRF DII, while the C-terminal region of DV of MsmEF-G2 is shorter and fails to form such interactions.

Notably, in the Msm50S–RRF–EF-G1 complex, domain IV plays a pivotal role in driving the rotation of RRF domain II toward the inter-subunit bridge B2a/d. The domain IV of MsmEF-G2 is displaced more towards the RRF domain II in a manner that would result in a near steric clash (Fig. 4B). Electrostatic surface potential analyses revealed that the interacting surface of MsmEF-G1 domain IV is optimally shaped and electrostatically complementary to MsmRRF domain II, whereas EF-G2 domain IV is not. (Fig. 4C and D). Further, in the Msm50S–RRF–EF-G1 complex, the C-terminal helix of MsmEF-G1 domain V extends towards MsmRRF domain I and makes connections to stabilise the RRF-ribosome interface. However, in EF-G2, this helix is notably shorter, precluding such contacts and thereby reducing its ability to stabilise RRF (Fig. 4E).

Extensive classification of our cryo-EM data captured another state in which MsmEF-G1 is bound to the 70S ribosome with tRNA, closely resembling a translocation state intermediate (Supplementary Fig. 6A). Comparison of this state with the 70S-P tRNA class shows that the 30S subunit adopts a ratcheted conformation (Supplementary Fig. 6B). The MsmEF-G1 occupies the inter-subunit space of the 70S ribosome. The MsmEF-G1 domain I (D1) is positioned in close proximity to the SRL, while the domain III (DIII) lies adjacent to the ribosomal protein bS12 on the 30S subunit (Supplementary Fig. 6E). The domain V (DV) of MsmEF-G1 interacts with the H43 of the 23S rRNA and bL11 (Supplementary Fig. 6F). The domain IV (DIV) of EF-G1 occupies the ribosomal A site on the 30S subunit in close proximity to the conserved nucleotide residues G510, A1476 and A1477 (*E.coli* numbering nt G530, A1492 and A1493) of the decoding centre (Supplementary Fig. 6G). This translocation intermediate state closely resembles the crystal structure reported by J.Zhou *et al*., where they have trapped *T.thermophilus* ribosomes bound with EF-G, mRNA, and tRNA in the presence of GDPNP(31). The overall conformation of the two EF-Gs is highly similar in both structures (Supplementary Fig. 6H), and in each case the tRNA occupies a hybrid position between the P and E site. However, in our structure, the ASL of the tRNA is displaced slightly further towards the E site (Supplementary Fig. 6D), suggesting that yet another intermediate state of tRNA has been trapped in the structure presented here. Structural analysis of this map indicated that EF-G1 is the translation factor responsible for carrying out tRNA translocation as well. Nevertheless, additional experiments are needed to confirm this claim.

Comparison of our Msm70S–EF-G1 structure with the previously reported Msm70S–EF-G2 structure (30) reveals pronounced differences in both the ribosome and EF-G conformations. In the Msm70S–EF-G2 complex, the 30S subunit adopts a non-ratcheted conformation, in contrast to the ratcheted state observed in our 70S–EF-G1 structure (Supplementary Fig. 6C). Previous studies have established that EF-G preferentially binds to and stabilises the ratcheted pre-translocation 70S ribosome during tRNA-mRNA translocation (6,46,47), however no such stabilisation is observed in the Msm70S-EF-G2 structure. We observe that MsmEF-G2 is shifted more towards the 30S subunit, with its domain I (D1) displaced away from the SRL, a key element required for efficient GTPase activation (Supplementary Fig. 6I). During the translocation process, the loops 1 and 2 of EF-G domain IV (DIV) make crucial contacts with the tRNA occupying a hybrid position between the A and the P site(6). These loops are correctly positioned in MsmEF-G1; in contrast, in MsmEF-G2, they are shifted away from the tRNA, abolishing these key interactions (Supplementary Fig. 6J).

Taken together, these observations provide a structural explanation for the functional divergence of mycobacterial EF-G isoforms. EF-G1 retains both ribosome-dependent GTPase activity and the structural compatibility required to remodel and stabilise RRF during ribosome recycling as well as during translocation. In contrast, MsmEF-G2 fails to show these properties and functions by trapping the post-translocated ribosome, thus slowing protein synthesis during stress(30).

### Conformational rearrangements in the 70S ribosome inter-subunit bridges upon RRF binding

The binding of RRF to the 70S ribosome stabilises the ratcheted conformation of the 30S subunit. In line with previous reports (48,49), this motion results in the rearrangement of multiple inter-subunit bridges that hold the two ribosomal subunits together(50). We aligned the non-ratcheted 70S-P tRNA map with Map 1 and analysed the changes in the inter-subunit bridges.

Bridge B1 was one of the first inter-subunit bridges identified, connecting the 30S head domain and the 50S central protuberance (CP). Bridge 1a (B1a) is formed between the ribosomal protein bS13 amino acid residues, Arg93 and Arg94 and the nucleotide residues nt A1003 and C1004 of H38 within the 23S rRNA. In Map 1, both amino acid residues Arg93 and Arg94 of the protein bS13 are shifted away from H38, thus disrupting the interactions (Supplementary Fig. 7A). Bridge B1b is formed between the ribosomal protein bS13 residues Val7, Asp8 and Gly68 with the residues Asp120, Val150 and Asp151 of the large subunit protein uL5. In Map 1, we observe that the positions of all the bS13 amino acid residues are altered, suggesting weakening of the bridge. Further change is observed in the interaction between the residues Leu4 and Arg57 of bS13 and Glu36 and Val37 of bL31, which also contributes to the formation of B1b (Supplementary Fig. 7B).

One of the most critical inter-subunit bridges is the Bridge B2, the pivotal point of ratchet rotation (48,50), which is subdivided into several interactions. Bridge B2a/d is situated close to the decoding centre and is formed between h44 nucleotide residues (nt 1476-1481) of the 30S ribosomal subunit and residues (nt 2135-2145) of H69 of the 50S ribosomal subunit, along with h45 residue G1501, and h24 residue U773, which interacts with C2144 and C2145 of H69, respectively. We observe local rearrangements in the interacting residues upon RRF binding (Supplementary Fig. 7C). Bridge B2b is formed through interactions between C763 of h24 and C2053 of H68, as well as between G1500 of h45 and U2155 of H71. We observe a shift in the h24 residue C763 due to the binding of RRF, resulting in the destabilisation of the bridge (Supplementary Fig. 7D). A similar observation is seen in the case of bridge B2c formed between C881, A882, A883 of h27 and C750 and G751 of h24 with C2049 and C2050 of H67, where the h24 residues are shifted away from H67, leading to the disruption of the interactions (Supplementary Fig. 7E).

Bridge B3 remains largely unaffected by the binding of RRF, which involves contacts between h44 residues (A1401, G1402, C1404, A1467 and U1468), H71 residues (nt 2171-2174 and 2182-2184) and uL14 residue (Lys53 and Arg54). Bridge B4 is formed by the interaction between the ribosomal protein uS15 and the loop region of H34. In Map 1, we observe that the interactions between the residues Gln37 and Arg88 of uS15 with A830 and A831 of H34 have noticeable changes (Supplementary Fig. 7F).

Another bridge where h44 is involved is the Bridge B5, where it interacts with the H62 of the 23S rRNA. Bridge B5 is formed between h44 (nt 1411, 1422, and 1459) and H62 (nt 1905, 1907, 1922 and 1918). Similar to bridge B3, this bridge shows minimal alteration upon RRF binding. In bridge B6, the h44 residue G1415 interacts with the L19 residue Lys105, and C1447 (h44) interacts with Lys108 (L19). Both G1415 and C1447 are shifted away from their original position, resulting in the weakening of the interactions (Supplementary Fig. 7G).

Bridge 7 is formed near the ribosomal E site, which links the base of the L1 stalk of the 50S subunit with the platform of the 30S subunit. In bridge B7a, the residue A682 (h23), which interacts with A2065 (H68), shows a considerable shift from its original position (Supplementary Fig. 7H). The bridge B7b is formed between h23 (U661 and A692) and h24 (G754 and G755) of the 16S rRNA and ribosomal protein uL2 residues (Leu165, Met177, Ser202 and Arg270). In Map 1, we observe that the h23 and h24 residues are shifted away from their position (Supplementary Fig. 7I). Bridge B8 connects h14 residues (338–341, 344–346) with uL14 residues (Asn13, Thr96, Arg97, and Glu121) and L19 residues (Ile39, Gln40, and Val41). The displacement of h14 residues in Map 1 leads to a disruption of the interaction (Supplementary Fig. 7J).

### Mechanistic pathway of RRF cooperating with EF-G1 to drive dissociation of ribosomal subunits

The binding of MsmEF-G1 to the RRF-bound 70S ribosome induces a pronounced ∼60° rotation of MsmRRF domain II (relative to its position in Map 1 and Map 3) towards the inter-subunit bridge B2a/d, resulting in its disruption and subsequent splitting of the 70S ribosome (Fig. 5A and B). Following subunit dissociation, the concerted action of MsmRRF domain II and MsmEF-G1 domain IV remodels the tip of H69 toward MsmRRF domain I (Map 4). This repositioning establishes new stabilising interactions between H69 and MsmRRF domain I. Specifically, the terminal nucleotide residues of H69 (C2138–C2139) form interactions with Lys12 and Lys15, while adjacent nucleotides A2140–U2141 interact with Ser127 and Asn130 of MsmRRF domain I. In addition, amino acid residues Ser47–Pro50 of the MsmRRF domain II establish contacts with nucleotide residues U2135–A2136 of H69, further consolidating this remodelled configuration (Fig. 5C).

**Figure. 5:**
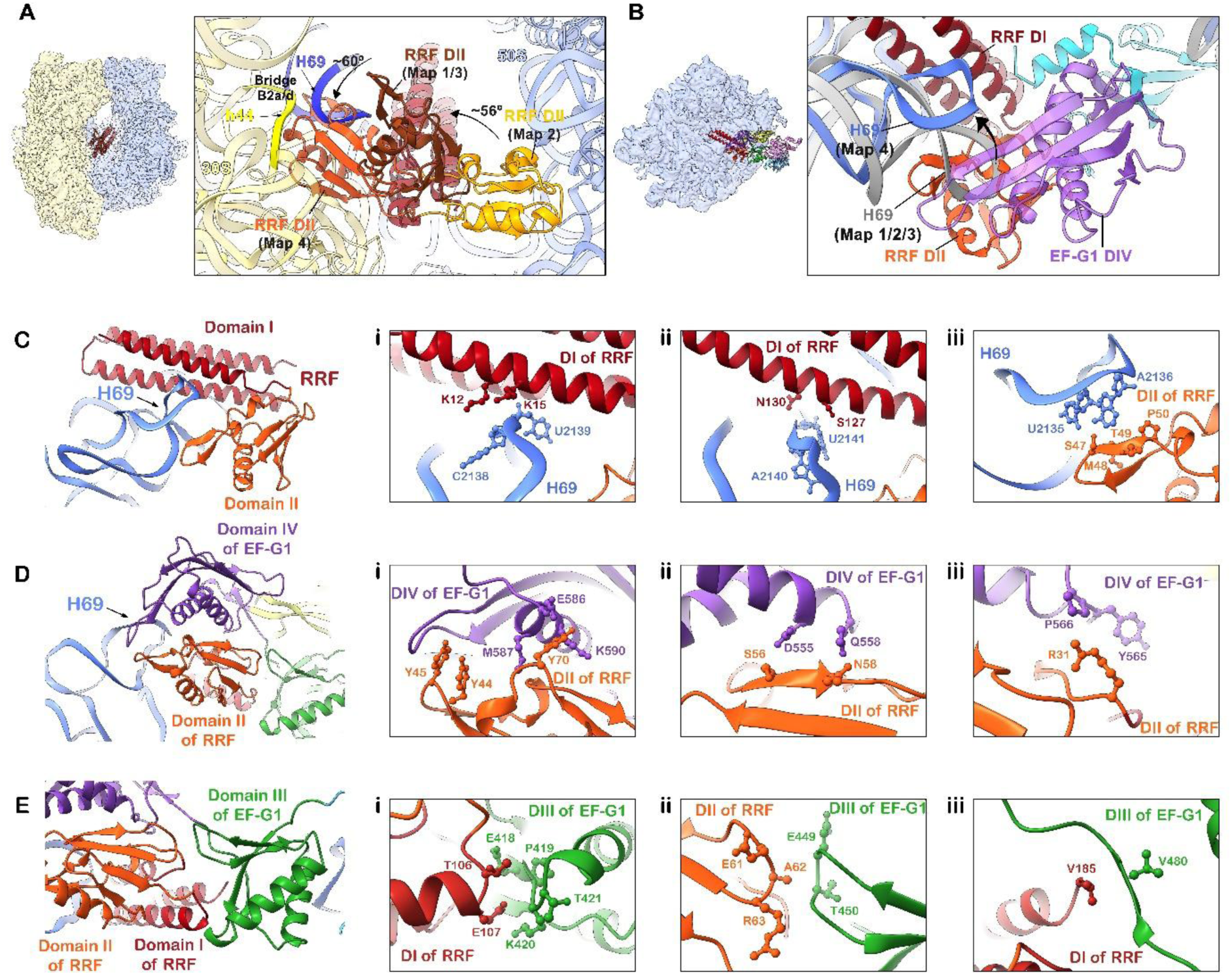
Conformational dynamics of MsmRRF and its coordinated interactions with MsmEF-G1 and H69 during ribosome recycling. (A) Close-up view showing the different conformation of MsmRRF domain II (DII) in Map 1/3 (red), Map 2 (mustard) and Map 4 (orange) across recycling intermediates. The MsmRRF DII makes a pronounced rotation of approximately 60° towards the inter-subunit bridge B2a/d in Map 4 when compared to Map 1/3. (B) The altered conformation of the H69 in Map 4. The MsmEF-G1 domain IV (DIV) (purple) pushes the MsmRRF DII (orange) towards the inter-subunit bridge B2a/d, leading to the shift in the position of H69 (blue) in Map 4 (blue) relative to its position in Maps 1/2/3 (grey). (C) Overview of the H69 (blue) interaction with MsmRRF domain I (DI) (red) (Ci,ii) and MsmRRF DII (orange) (Ciii). (D) Overview of the MsmRRF DII (orange) interaction with MsmEF-G1 DIV (purple). (E) Overview of the interaction of MsmEF-G1 domain III (DIII) (green) with MsmRRF DI (red) (Ei, iii) and MsmRRF DII (orange) (Eii).

The MsmEF-G1-assisted repositioning of domain II of MsmRRF establishes a network of new interactions between MsmRRF and the 23S rRNA. Although domain IV of MsmEF-G1 does not directly contact helix H69, it plays a critical role in stabilising the rotated MsmRRF domain II through multiple inter-domain interactions. Specifically, Tyr44 and Tyr45 of MsmRRF domain II interact with loop I residues of MsmEF-G1 domain IV, whereas Tyr70 of MsmRRF domain II contacts residues Glu586-Lys590 of MsmEF-G1 domain IV. Furthermore, Ser56 and Asn58 of MsmRRF domain II form hydrogen bonds with Asp555 and Gln558 of MsmEF-G1 domain IV, respectively. The linker residue Arg31 of MsmRRF also interacts with Tyr565 and Pro566 of MsmEF-G1 domain IV, thereby reinforcing the interface between the two factors (Fig. 5D)

Additionally, domain III of EF-G1 establishes auxiliary contacts with domains I and II of MsmRRF, further stabilising the complex. Notably, Thr106 and Glu107 of MsmRRF domain II interact with the Glu418-Thr421 stretch of MsmEF-G1 domain III, while Glu61-Arg63 of MsmRRF domain II form electrostatic interactions with Glu449-Thr450 of MsmEF-G1 domain III. Moreover, the hydrophobic interaction of Val185 of MsmRRF domain I occurs with Val480 and Gly481 of MsmEF-G1, providing the interfacial stability (Fig. 5E).

Altogether, these interactions stabilise the rotated conformation of MsmRRF domain II and reinforce its engagement with H69. The stabilisation extends to adjacent helix H71, where residues (nt C2166, U2167, G2169, U2187 and C2189) interact with MsmRRF domain I residues Lys125, Val126, Arg129 and Arg133, thereby stabilising the post-dissociation state of the ribosome. These findings establish a structural framework in which MsmEF-G1 facilitates ribosome splitting by inducing domain II rotation of MsmRRF and subsequently consolidates the remodelled MsmRRF and ribosome interaction to ensure efficient recycling.

Based on the different structural states obtained in this study, we propose sequential steps for the ribosome recycling process in mycobacteria. The post-termination complex (PoTC) carrying a deacylated tRNA at the P site represents an ‘unlocked’ state of the ribosome (49) where the PoTC interconverts between the ratcheted and the non-ratcheted state spontaneously (Fig. 6, Step I). RRF approaches and binds to PoTC in the most stable conformation, as represented by the isolated RRF crystal structure (Supplementary Fig. 8B) and stabilises the ratcheted state of the ribosome (51,52). Surface potential of MsmRRF domain I indicates that a repulsive interaction with the tRNA CCA end, in addition to steric collision, aids in pushing the tRNA into the P/E hybrid state, stabilising the ratcheted state of the 30S subunit (Supplementary Fig. 8A). In this ribosome-bound state, RRF domain I extends towards the P site on the 50S ribosomal subunit and domain II is oriented towards the 30S subunit and interacts with bS12 (Fig 6. Step II) (Map 1). The domain I forms multiple contacts with ribosomal components, such as H69, H71, H80, and protein uL16, which contribute to its stability on the ribosome. The tRNA occupies a P/E hybrid position, and the 30S subunit has a ratchet-like motion which promotes the rearrangement of multiple inter-subunit bridges. Notably, RRF alone cannot induce ratchet motion as shown by the non-ratcheted 70S-RRF complex, suggesting that the presence of deacylated tRNA at the P site is needed (Supplementary Fig. 4B). While the head of the 30S subunit remains in a swivelled position, the body rotates back to its original state, positioning the tRNA in the E/E position (Maps 2 and 3). The domain II of RRF is flexible and is seen to oscillate between the 50S and the 30S subunits (Fig.6, Steps IIIa and b). The subsequent binding of EF-G1 orients RRF domain II further towards the inter-subunit bridge B2a/d, causing its disruption instantaneously, leading to the dissociation of the 70S ribosome, and the release of mRNA and the tRNA.

**Figure. 6:**
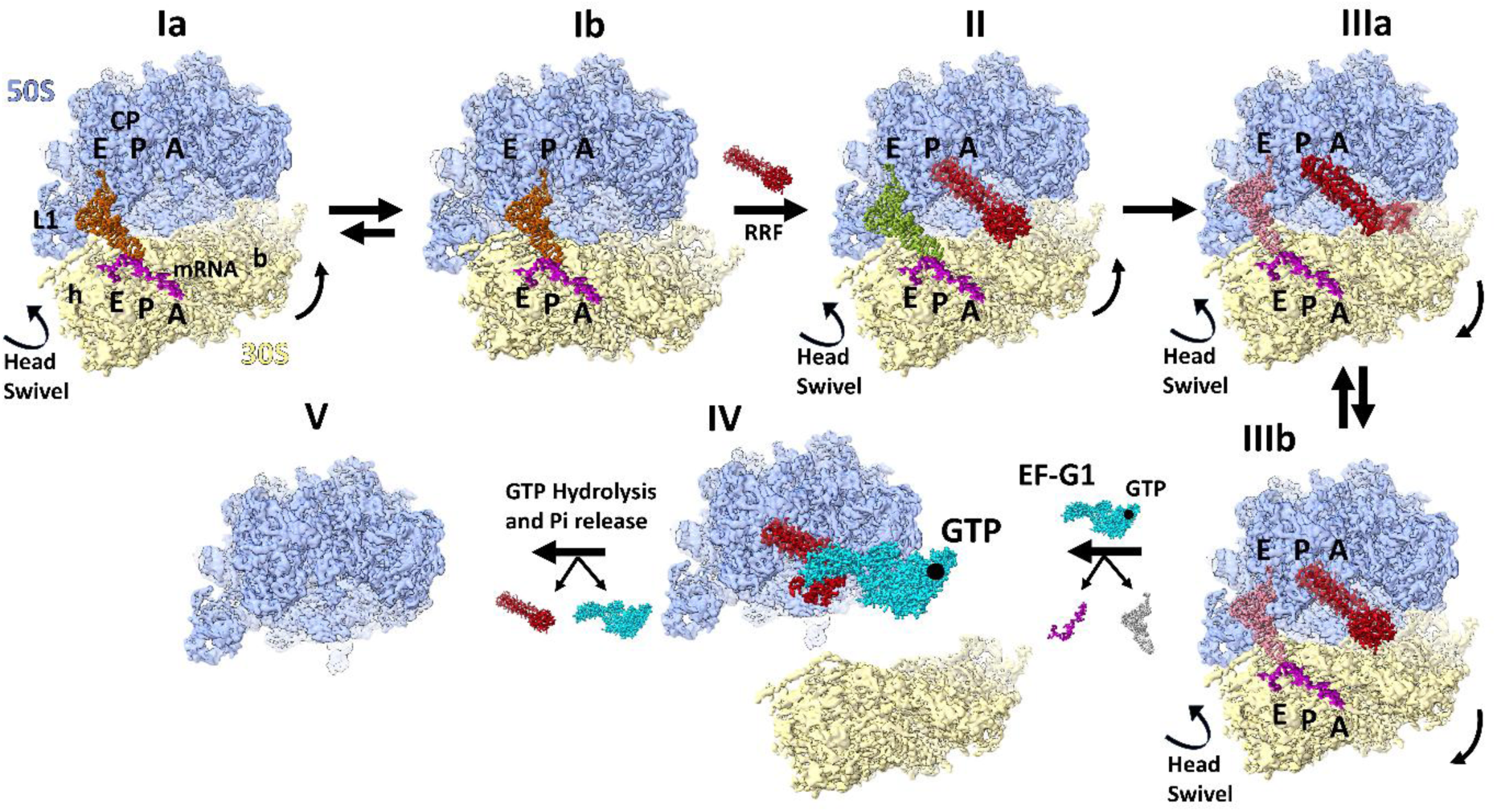
Schematic representation of the proposed model of MsmRRF and MsmEF-G1-mediated ribosome recycling process in Mycobacterium. State Ia–Ib depict the post-termination 70S ribosome (PoTC) containing a deacylated tRNA (orange) at the P site and mRNA (magenta). The PoTC interconverts between the ratcheted (Ia) and the non-ratcheted state (Ib). State II shows binding of MsmRRF (red) to the 70S ribosome with its domain II oriented towards the 30S subunit (yellow). This binding of MsmRRF stabilises the ribosome in the ratcheted conformation with the tRNA occupying the P/E hybrid state (green). State IIIa-IIIb represents the state where the body of the 30S subunit rotates back to its original position while the head remains in the swivelled position, placing the tRNA in the E/E position (light pink). The MsmRRF domain II (DII) oscillates between the 50S (IIIa) and the 30S subunits (IIIb). State IV represents the recruitment of the MsmEF-G1 (cyan) to the RRF-bound ribosome, where the domain IV (DIV) of MsmEF-G1 orients the MsmRRF DII towards the inter-subunit bridge B2a/d, causing its disruption. This leads to the dissociation of the 70S ribosome and the subsequent release of the mRNA and the tRNA. The MsmRRF and the MsmEF-G1 remain bound to the 50S subunit (blue). State V shows that the MsmRRF and the MsmEF-G1 are released from the 50S subunit upon GTP hydrolysis. CP-central protuberance, h-head, b-body.

Earlier studies have shown that in bacteria, the dissociation of the 70S ribosome requires GTP hydrolysis (8,9,18,53). However, reports show that in *B. burgdorferi* and human mitochondria, the process is GTP-independent (26,28). In mitochondria, the GTP hydrolysis is required to release the mtEF-G2 and mtRRF from the large ribosomal subunit (28). In agreement with observations in human mitochondria, our structural data show that in the presence of the non-hydrolysable GTP analogue (GMP-PNP), the mycobacterial EF-G1 and RRF remain bound to the 50S subunit after the 70S dissociation (Map 4). Supporting this observation, the 70S ribosome dissociation assay performed in the presence of GMP-PNP showed that the 70S ribosome was able to split into its constituent subunits (Supplementary Fig. 5A-C). So, as observed in the case of mitochondrial and *B. burgdorferi* ribosome recycling, 70S dissociation is GTP-independent and is required only to release EF-G1 and RRF from the 50S subunit (Fig.6, Steps IV and V). However, further experiments are required to validate this claim.

## Conclusion

In this study, we report for the first time a comprehensive set of high-resolution structures showcasing multiple intermediates of the mycobacterial ribosome recycling process. All the reconstructed maps reach global resolutions in the range of approximately 3-4 Å. We were unable to visualise the transient intermediate in which both MsmRRF and MsmEF-G1 are simultaneously bound to the 70S ribosome, immediately preceding subunit dissociation. As reported, this process occurs extremely fast, thus making this complex particularly challenging to trap structurally(8).

The 30S subunit also exhibits distinct conformational states, including a ratcheted conformation in Map 1 and pronounced head swivelling in Maps 2 and 3. The binding of MsmRRF to the ribosome helps to stabilise the ribosome in the ratcheted state. The high resolution of our maps enabled a detailed examination of inter-subunit bridges, many of which undergo substantial rearrangements occurring due to this ratchet-like motion of the 30S subunit.

Our study establishes MsmEF-G1 as the structurally and functionally competent factor that, together with MsmRRF, mediates ribosome recycling in mycobacteria. The MsmRRF domain I is very stable and occupies the same position in all the maps, in contrast to some of the previous reports (18,54), anchors through extensive interactions with helices H69, H71, and H80 of the 23S rRNA and ribosomal protein uL16. The binding of MsmEF-G1 to the Msm70S-RRF complex induces a pronounced rotation of MsmRRF domain II about its hinge region toward the inter-subunit bridge B2a/d, leading to the disruption of this critical bridge, thereby promoting the splitting of the 70S ribosome. Detailed structural analysis of the G domain of MsmEF-G1 and mtEF-G2 revealed that the G Motifs are well conserved and thus shows ribosome dependent GTPase activity. In addition, MsmEF-G1 has a compatible electrostatic surface potential as well as shape features necessary to establish a stable and productive interface with MsmRRF. In contrast, the key amino acid residues of the G domain in MsmEF-G2 are substituted, and the G domain itself is shifted away from the SRL. It lacks ribosome-dependent GTPase activity and has unfavourable electrostatic interactions with the MsmRRF, rendering it inactive for ribosome recycling. Consistent with the observations in mitochondrial and *B. burgdorferi* ribosome recycling, our structural as well as biochemical analyses indicate that in mycobacteria as well, the disassembly of the 70S ribosome is independent of GTP hydrolysis. Rather, GTP hydrolysis is mainly required for the subsequent release of the associated protein factors from the large ribosomal subunit.

The structural characterisation of a Msm70S-EF-G1 complex corresponding to a translocation intermediate indicates that MsmEF-G1 is the canonical translation factor involved not only in ribosome recycling but also in the tRNA translocation step.

## Acknowledgements.

This work was primarily supported by SERB, DST (India) sponsored project (SPF/2021/000141). We also acknowledge funding from CSIR-Indian Institute of Chemical Biology (CSIR-IICB), Kolkata, India (OLP-120). We gratefully acknowledge the National Electron Cryo microscopy facility at the Bangalore Life Sciences Cluster (DBT/PR12422/MED/31/287/2014), NCBS, Bangalore, DBT (grant no. BT/PR15017/BRB/10/1445), and the Central Instrumentation Facility (CIF) of CSIR-IICB. We sincerely thank Dr. Vinothkumar Kutti Ragunath and Dr. Sucharita Bose for helping in grid preparation and high-resolution single-particle data collection. AD, PB, KS acknowledge UGC and CSIR for the research fellowships.

## Author Contribution

JS conceived the project. AD and PB designed and carried out all the experiments in consultation with JS. AD and KS did the cryo-EM 3D data-processing. JS and AD analysed the data, wrote the paper and prepared the illustrations.

## Availability of data

For Map 1, Map 2, Map 3, Map 4, 70S-RRF, 70S-EFG1 and 70S-PtRNA, the cryo-EM density maps have been deposited in the Electron Microscopy Data Bank (EMDB) under IDs EMD-69697, EMD-69698, EMD-69703, EMD-69705, EMD-69707, EMD-69709, and EMD-69708, respectively. Coordinates are deposited in the Protein Data Bank (PDB) under IDs 24NM, 24NN, 24NP, 24NS, 24NU, 24NW and 24NV, respectively.

---

**Table S1.** Data collection, refinement and validation statistics.

| Ribosome Complex | Map 1 | Map 2 | Map 3 | Map4 | 70S-RRF | 70S-EFG1 | 70S-PtRNA |
| --- | --- | --- | --- | --- | --- | --- | --- |
| Database submission |  |  |  |  |  |  |  |
| EMDB ID | EMD-69697 | EMD-69698 | EMD-69703 | EMD-69705 | EMD-69707 | EMD-69709 | EMD-69708 |
| PDB ID | 24NM | 24NN | 24NP | 24NS | 24NU | 24NW | 24NV |
| Data Collection |  |  |  |  |  |  |  |
| Microscope | Titan Krios |  |  |  |  |  |  |
| Camera | Falcon III (4k x 4k) |  |  |  |  |  |  |
| Nominal Magnification | 75000x |  |  |  |  |  |  |
| Voltage (keV) | 300 |  |  |  |  |  |  |
| Electron Dose (e/ Å²) | 20.82 |  |  |  |  |  |  |
| Defocus Range (um) | -1.8 to -3.3 |  |  |  |  |  |  |
| Pixel Size (Å) | 1.07 |  |  |  |  |  |  |
| Cryo-EM Reconstruction |  |  |  |  |  |  |  |
| Refined Map Resolution (Å) | 4.5 Å | 5.1 Å | 4.2 Å | 6.6 Å | 3.7 Å | 5.1 Å | 3.9 Å |
| Postprocessed Map Resolution(Å) | 3.6 Å | 3.8 Å | 3.4 Å | 4.4 Å | 3.1 Å | 3.7 Å | 3.2 Å |
| Final Particle Number | 24,608 | 18,050 | 32,720 | 15,686 | 61,327 | 13,826 | 36,773 |
| Symmetry | C1 |  |  |  |  |  |  |
| FSC Threshold | 0.143 |  |  |  |  |  |  |
| Resolution Metric | FSC |  |  |  |  |  |  |
| Model Refinement |  |  |  |  |  |  |  |
| Model Resolution (Å) | 3.2 Å | 3 Å | 3 Å | 3.5 Å | 2.8 Å | 3.2 Å | 2.9 Å |
| FSC Threshold | 0.143 |  |  |  |  |  |  |
| Map-Sharpening B factor (Å²) | -122.1 | -132.4 | -122.3 | -177.8 | -120.4 | -136.3 | -111.8 |
| Model Composition |  |  |  |  |  |  |  |
| Chains | 71 | 70 | 70 | 47 | 69 | 71 | 70 |
| Non-hydrogen Atoms | 152926 | 151061 | 151086 | 104614 | 149587 | 155100 | 149778 |
| Protein Residues | 6464 | 6230 | 6233 | 4567 | 6226 | 6749 | 6045 |
| Nucleotide Residues | 4794 | 4793 | 4793 | 3237 | 4721 | 4793 | 4797 |
| Ligands | ZN:6<br>MG:12 | ZN:6<br>MG:12 | ZN:6<br>MG:12 | GNP:1<br>ZN:4<br>MG:10 | ZN:6<br>MG:12 | GNP:1<br>ZN:6<br>MG:12 | ZN:6<br>MG:12 |
| Root-mean-square Deviations |  |  |  |  |  |  |  |
| Bond Length (Å) | 0.005 | 0.008 | 0.008 | 0.008 | 0.007 | 0.003 | 0.006 |
| Bond Angle (°) | 0.845 | 0.991 | 0.988 | 0.853 | 0.719 | 0.601 | 0.680 |
| Validation |  |  |  |  |  |  |  |
| MolProbity Score | 1.83 | 1.76 | 1.75 | 1.63 | 1.83 | 1.84 | 1.80 |
| Clash Score | 9.60 | 9.0 | 8.85 | 6.79 | 8.73 | 10.44 | 8.25 |
| Rotamer Outliers(%) | 0.69 | 0.46 | 0.46 | 1.03 | 0.1 | 0.24 | 0.02 |
| Ramachandran Plot |  |  |  |  |  |  |  |
| Outliers (%) | 0 | 0.02 | 0.02 | 0.02 | 0.07 | 0 | 0.05 |
| Allowed (%) | 4.66 | 4.05 | 4.05 | 3.67 | 5.10 | 4.31 | 5.12 |
| Favoured (%) | 95.34 | 95.93 | 95.94 | 96.31 | 94.84 | 95.69 | 94.83 |

