## Supplementary Material for "Structural investigation suggests Elongation Factor-G1 is the *bona fide* canonical translation factor for ribosome recycling in mycobacteria"

Supplementary Data

**Supplementary Figure. 1:** Comparative sequence analysis of MsmRRF and MsmEF-Gs across different species. (A) Multiple sequence alignment (MSA) of MsmRRF from *Mycobacterium smegmatis*, *Mycobacterium tuberculosis*, *Mycobacterium avium*, *Mycobacterium ulcerans, Escherichia coli*, *Thermus thermophilus*, *Borrelia burgdorferi*, and human mitochondria. The MSA was generated using Clustal Omega. (B) Multiple sequence alignment (MSA) of the G domain of EF-Gs from *Escherichia coli, Mycobacterium smegmatis*, *Mycobacterium tuberculosis, Borrelia burgdorferi*, and human mitochondria. (C-F) Percentage identity matrix generated using Clustal Omega showing the percentage of amino acid sequence identity of MsmRRF across the above-mentioned species (C). Percentage sequence identity of MsmEF-G1 (D) and MsmEF-G2 (E) between *Escherichia coli, Mycobacterium smegmatis* and *Mycobacterium tuberculosis* and of MsmEF-G1 and MsmEF-G2 between *Mycobacterium smegmatis* and *Mycobacterium tuberculosis* (F).

**Supplementary Figure. 2:** Schematic representation of the Cryo-EM single particle 3D reconstruction workflow. The good 2D class averages were selected, and they were subjected to multiple rounds of reference-based 3D classification followed by focused 3D classification. We obtained four high-resolution ribosome recycling intermediates, Map 1 (4.5Å), Map 2 (5.1Å), Map 3 (4.2Å) and Map 4 (6.5Å). In addition, we also obtained 70S-RRF (3.7Å), 70S-P tRNA (3.9Å) and 70S-EF-G1 map (5.1Å).

**Supplementary Figure. 3:** Cryo-EM single particle 3D reconstruction, local resolution estimation and FSC calculation. (A-G) The local resolution plot of the refined maps generated using RELION 3.1, the angular distribution of the particles and the Fourier Shell Correlation (FSC) plot showing the resolution of the refined map, where the resolution value is given according to “gold standard” (FSC=0.143) criteria. Map 1 (A), Map 2 (B), Map 3 (C), 70S-P tRNA (D), 70S-RRF (E), 70S-EF-G1 (F) and Map 4 (G).

**Supplementary Figure. 4:** Non-ratcheted conformation of the 30S subunit in 70S-RRF map and the analysis of the position of H43 in Map 2. (A) Cryo-EM map showing MsmRRF (red) bound to the 70S ribosome (30S subunit non-ratcheted) with a resolution of 3.1 Å. The 30S subunit is shown in yellow, and the 50S subunit is shown in blue. (B) Superposition of the 30S subunits in the structures 70S-P tRNA (purple) and 70S-RRF (cyan), showing the non-ratcheted 30S subunit. (C) Comparison of the position of H43 in Map 2 (blue), 70S-P tRNA class (cyan) and PDB 3JOD (grey) shows that H43 is shifted in the model PDB 3JOD.

**Supplementary Figure. 5:** Ribosome dissociation and GTPase assay. (A-C) The ribosome splitting profiles of Msm ribosome with MsmRRF, Msm EF-G1, *E.coli* IF3 in the presence of GTP (A), GMP-PNP (B) and no nucleotide (C). The ribosome profiles clearly show the dissociation of the 70S ribosome into the 50S and 30S subunits in the presence of GTP and GMP-PNP. (D) Malachite green assay showing the ribosome-dependent GTP hydrolysis by wild-type EF-G1, in contrast to the EF-G1 mutants R62Q and H88Y, showing reduced and near complete loss of GTPase activity, respectively.

**Supplementary Figure. 6:** 70S ribosome-bound EF-G1 represents a translocation intermediate state. (A) Cryo-EM map showing MsmEF-G1 (cyan) bound to the 70S ribosome with a resolution of 3.7 Å. The 30S subunit is shown in yellow, and the 50S subunit is shown in blue. (B) The conformational dynamics of the ribosomal 30S subunit. Superposition of the 30S subunits in the structures 70S-P tRNA (purple) and 70S-EF-G1(light green), showing the ratchet motion of the 30S subunit. (C) Superposition of the 30S subunits in the structures 70S-EF-G1 (light green) and 70S-EF-G2 (magenta), showing the ratchet motion of the 30S subunit. (D) Superposition of the tRNA in the structures 70S-EF-G1 (purple) and PDB 4V9J (grey), showing its conformational dynamics. The P/P tRNA (light blue) from the 70S-P tRNA structure has been shown as a reference. (E-G) Close-up views showing the position of binding of EF-G1 on the 70S ribosome, where MsmEF-G1 domain III (DIII) (light green) is in close proximity to bS12 (blue) (E), the MsmEF-G1 domain V (DV) (yellow) is in close proximity to H43 (blue) and bL11 (light brown) (F) and the MsmEF-G1 domain IV (DIV) (purple) occupies the A site and is in close proximity to the residues G510, A1476 and A1477 (orange) of the decoding centre (DC) (G). (H) Comparison of the conformation of MsmEF-G1 in the 70S-EF-G1 map (cyan) with the structure PDB 4V9J (grey). (I) Comparison of the conformation of MsmEF-G1 in the 70S-EF-G1 map (cyan) with MsmEF-G2 in the structure PDB 9K0Z (grey), with the sarcin-ricin loop (SRL) shown in blue, showing that MsmEF-G2 is shifted towards the 30S subunit away from the SRL. (J) Comparison of the conformation of MsmEF-G1 domain IV (DIV) in the 70S-EF-G1 map (cyan) with MsmEF-G2 domain IV (DIV) in the structure PDB 9K0Z (grey), showing that the loop 1 and 2 of MsmEF-G2 DIV is shifted away from the ap/P tRNA (grey) (PDB 7SSD).

**Supplementary Figure. 7:** Structural remodelling of the inter-subunit bridges during ribosome recycling. (A-J) Comparison of the 70S-P tRNA model (orange) with the Map 1 model (yellow), showing the residues of 70S-P tRNA involved in the formation of the inter-subunit bridge in blue and the altered positions of the corresponding residues of Map 1 in cyan, in Bridge B1a (A), Bridge B1b (B), Bridge B2a/d (C), Bridge B2b (D), Bridge B2c (E), Bridge 4 (F), Bridge B6 (G), Bridge B7a (H), Bridge B7b (I), Bridge B8 (J).

**Supplementary Figure. 8:** The domain architecture of RRF. (A) Electrostatic surface representation of MsmRRF, highlighting the negatively charged tip of the RRF domain I, which will both electrostatically repel the negatively charged CCA end of the P/P tRNA (cyan) as well as cause a steric clash. (B) Superposition of the previously reported isolated crystal structures of RRF (PDB 1WQH, 1WQG, and 4KDD) in the MsmRRF structure from Map 1 and Map 2 shows that the RRF domain II in all the crystal structures aligns closely with the of Map 1 conformation (red), oriented towards the 30S subunit, whereas in Map 2 the domain II (mustard) has a orientation towards the 50S subunit.

A

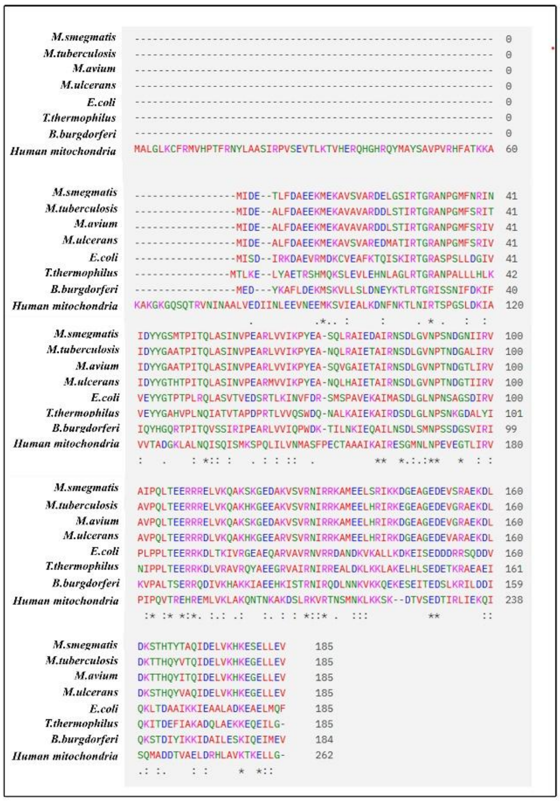


B

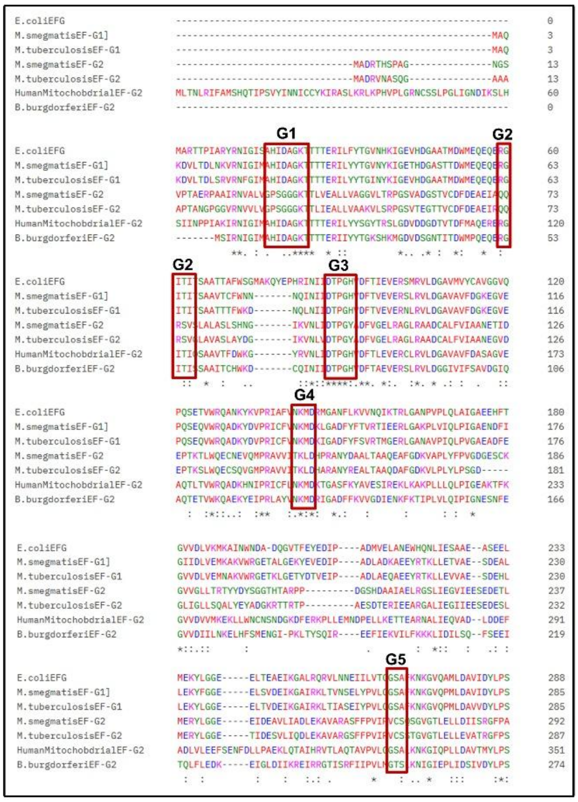


C

| | | | | | | | | |
|---|---|---|---|---|---|---|---|---|
| *M.smegmatis* | 100.00 | 85.95 | 87.03 | 85.41 | 40.54 | 39.13 | 44.02 | 28.02 |
| *M.tuberculosis* | 85.95 | 100.00 | 95.14 | 90.81 | 40.00 | 39.67 | 44.57 | 27.47 |
| *M.avium* | 87.03 | 95.14 | 100.00 | 89.73 | 42.16 | 38.04 | 43.48 | 27.47 |
| *M.ulcerans* | 85.41 | 90.81 | 89.73 | 100.00 | 42.16 | 40.76 | 44.02 | 26.37 |
| *E.coli* | 40.54 | 40.00 | 42.16 | 42.16 | 100.00 | 44.02 | 38.59 | 28.02 |
| *T.thermophilus* | 39.13 | 39.67 | 38.04 | 40.76 | 44.02 | 100.00 | 38.25 | 27.87 |
| *B.burgdorferi* | 44.02 | 44.57 | 43.48 | 44.02 | 38.59 | 38.25 | 100.00 | 25.41 |
| *Human mitochondria* | 28.02 | 27.47 | 27.47 | 26.37 | 28.02 | 27.87 | 25.41 | 100.00 |

D

| | | | |
|---|---|---|---|
| *E.coli* | 100.00 | 60.06 | 59.20 |
| *M.smegmatis* | 60.06 | 100.00 | 87.45 |
| *M.tuberculosis* | 59.20 | 87.45 | 100.00 |

E

| | | | |
|---|---|---|---|
| *E.coli* | 100.00 | 31.40 | 32.60 |
| *M.smegmatis* | 31.40 | 100.00 | 78.12 |
| *M.tuberculosis* | 32.60 | 78.12 | 100.00 |

F

| | | |
|---|---|---|
| *M.Smegmatis* EF-G1 | 100.00 | 31.25 |
| *M.Tuberculosis* EF-G2 | 31.25 | 100.00 |

Supplementary Figure. 1.

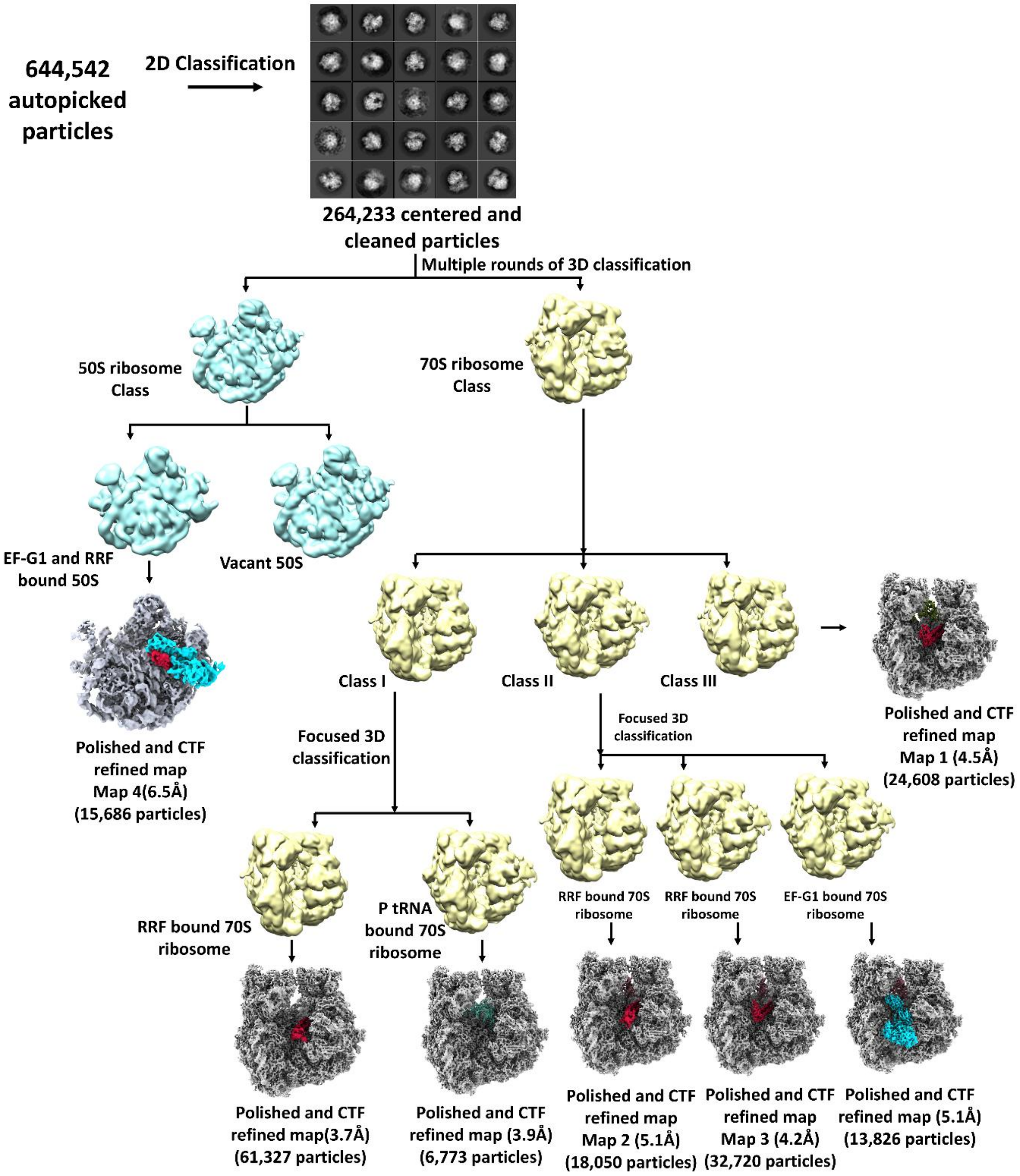


Supplementary Figure. 2.

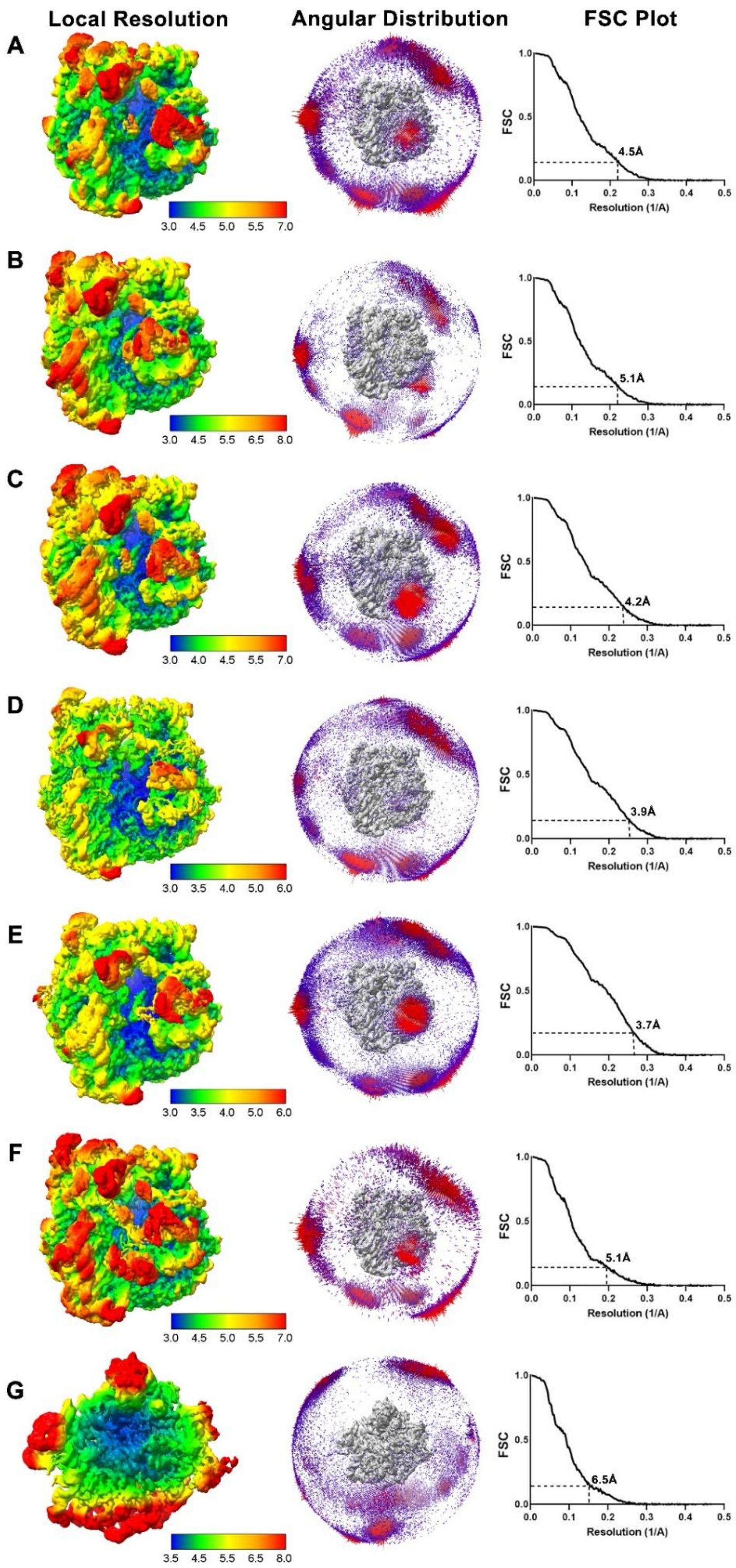


Supplementary Figure. 3.

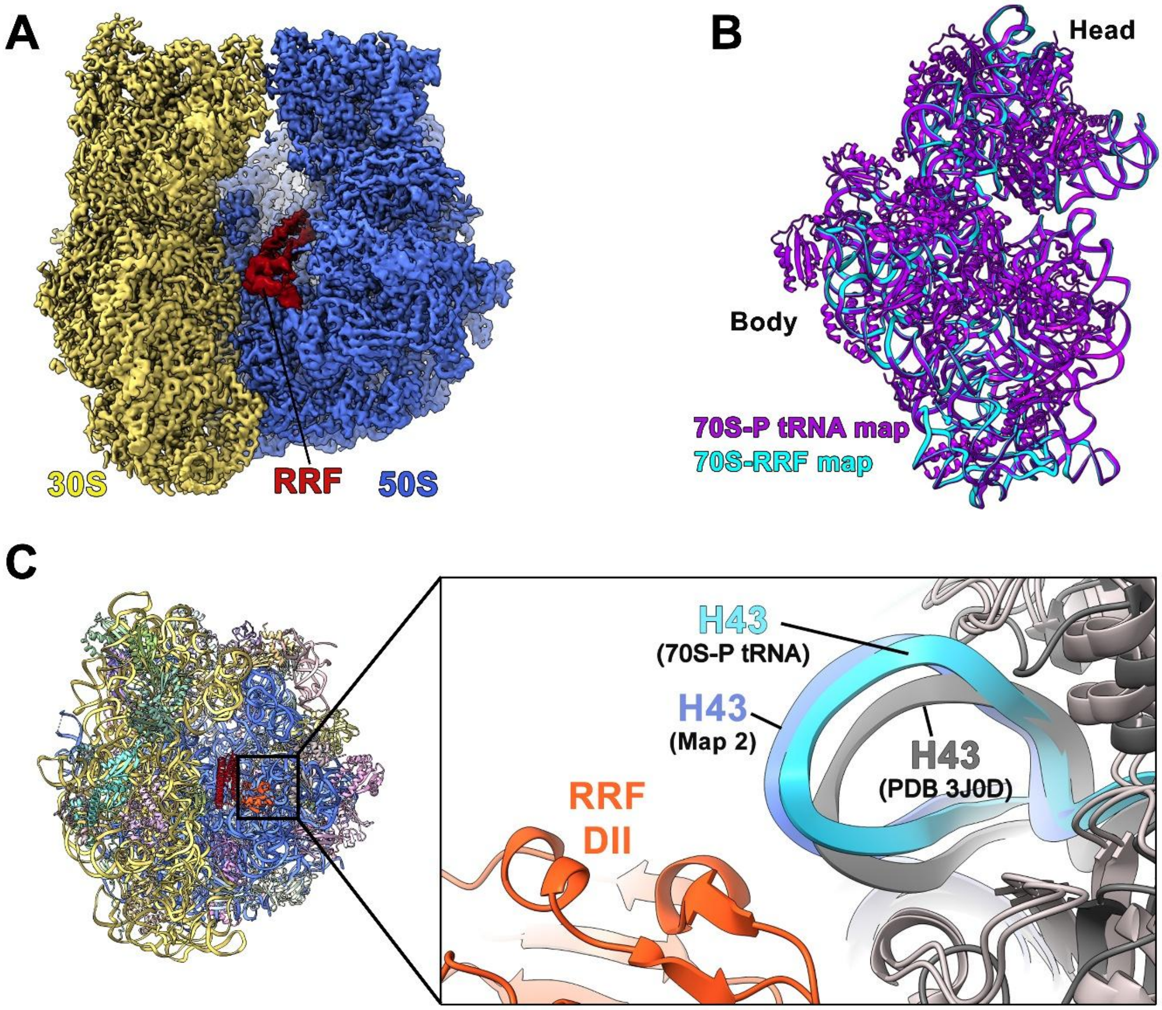


Supplementary Figure. 4.

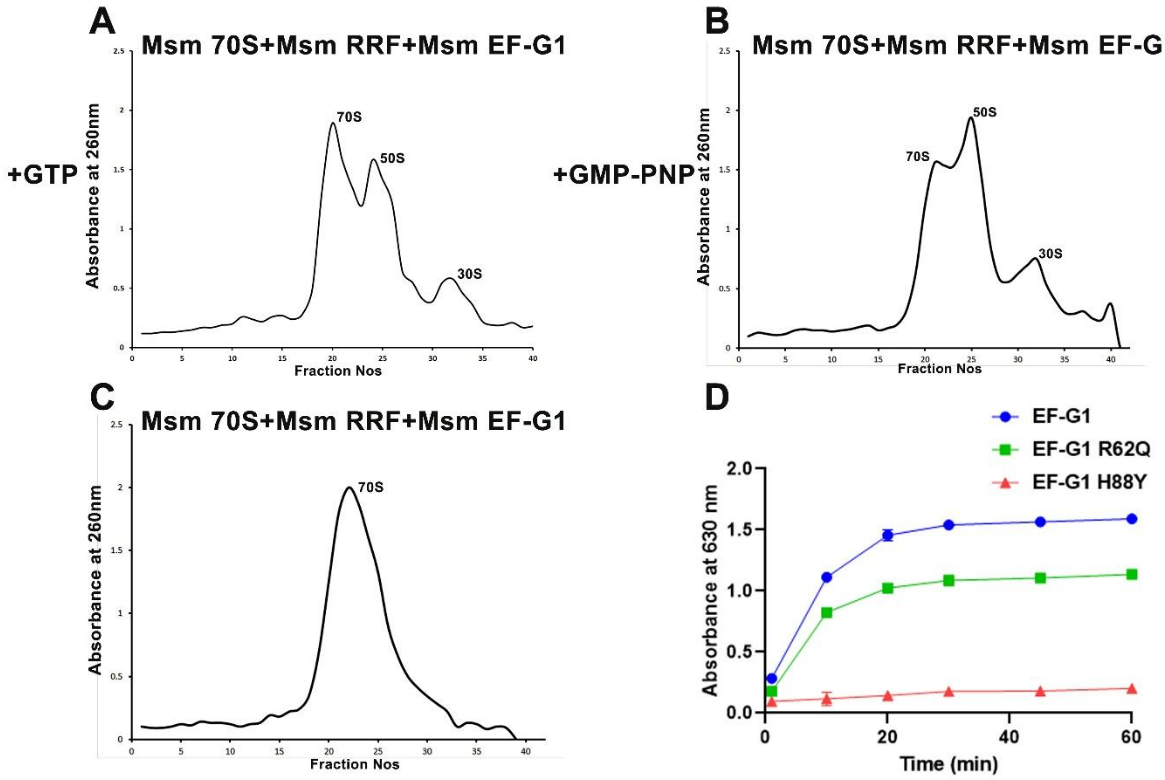


Supplementary Figure. 5.

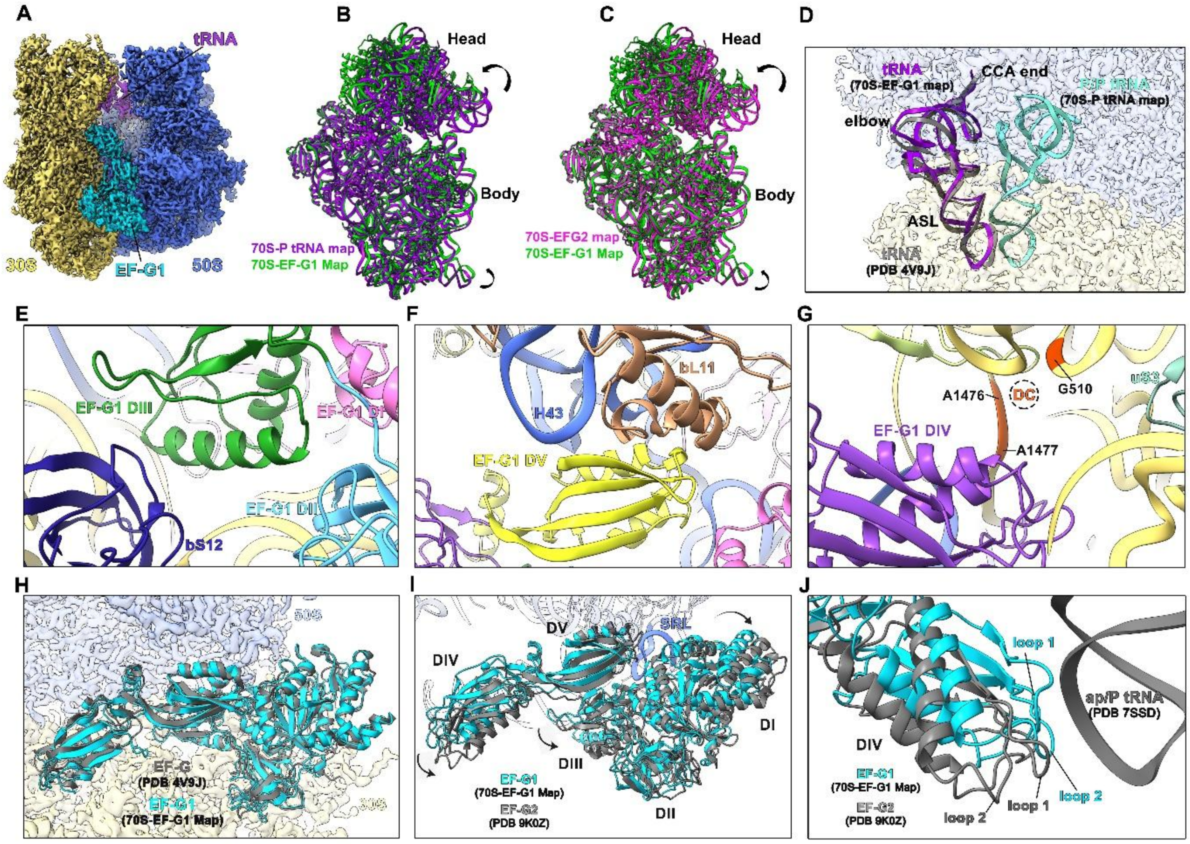


Supplementary Figure. 6.

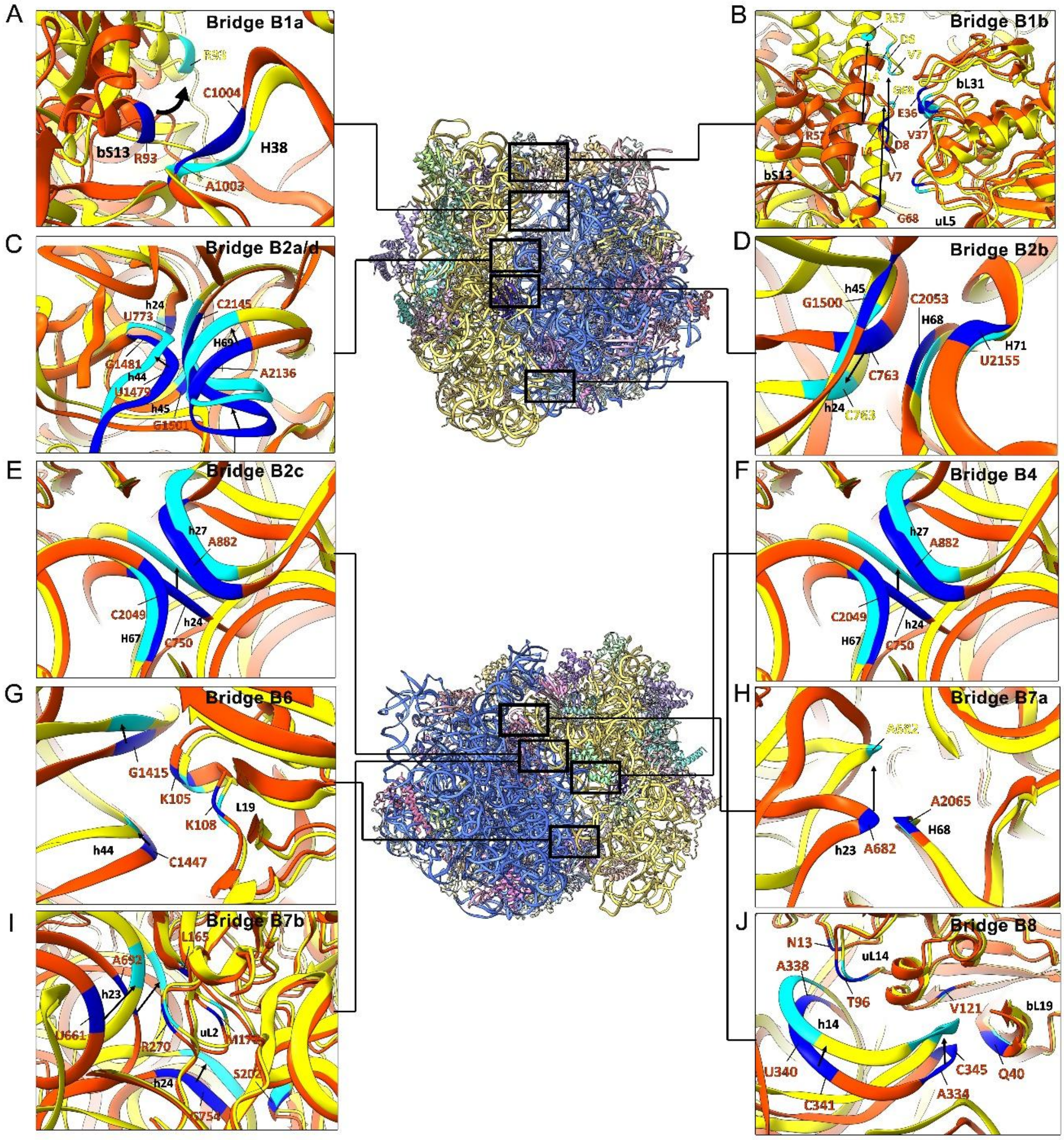


Supplementary Figure. 7.

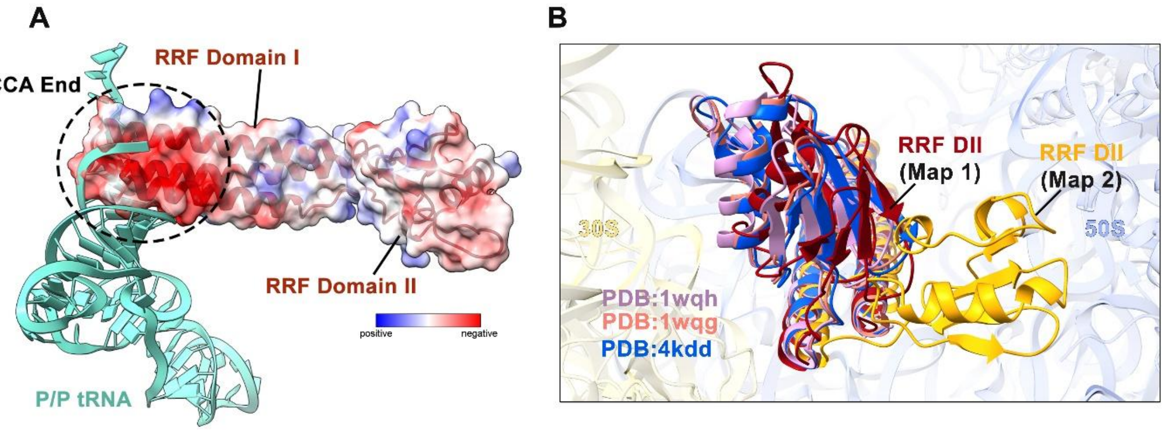


Supplementary Figure. 8.
